# Another’s pain in my brain: No evidence that placebo analgesia affects the sensory-discriminative component in empathy for pain

**DOI:** 10.1101/2020.05.18.101238

**Authors:** Helena Hartmann, Markus Rütgen, Federica Riva, Claus Lamm

## Abstract

The shared representations account of empathy suggests that sharing other people’s emotions relies on neural processes similar to those engaged when directly experiencing such emotions. Recent research corroborated this by showing that placebo analgesia resulted in reduced pain empathy and decreased activation in shared neural networks. However, those studies did not report any placebo-related variation of somatosensory engagement during pain empathy. The experimental paradigms used in these studies did not direct attention towards a specific body part in pain, which may explain the absence of effects for somatosensation. The main objective of this preregistered study was to implement a paradigm overcoming this limitation, and to investigate whether placebo analgesia may also modulate the sensory-discriminative component of empathy for pain. We induced a localized, first-hand placebo analgesia effect in the right hand of 45 participants by means of a placebo gel and conditioning techniques, and compared this to the left hand as a control condition. Participants underwent a pain task in the MRI scanner, receiving painful or non-painful electrical stimulation on their left or right hand, or witnessing another person receiving such stimulation. In contrast to a robust localized placebo analgesia effect for self-experienced pain, the empathy condition showed no differences between the two hands, neither for behavioral nor neural responses. We thus report no evidence for somatosensory sharing in empathy, while replicating previous studies showing overlapping brain activity in the affective-motivational component for first-hand and empathy for pain. Hence, in a more rigorous test aiming to overcome limitations of previous work, we again find no causal evidence for the engagement of somatosensory sharing in empathy. Our study refines the understanding of the neural underpinnings of empathy for pain, and the use of placebo analgesia in investigating such models.

**Highlights:**

- Investigated placebo modulation of somatosensory and affective components of pain
- Localized placebo analgesia effects for self-report and fMRI of first-hand pain
- No evidence for such effects in empathy for pain
- Suggests that somatosensory sharing does not play a critical role in pain empathy

## 1 Introduction

Empathy is a multifaceted psychological construct fundamental for human social interactions and relationships (e.g. Marsh, 2018 for recent review). While many definitions of empathy have been proposed, here we define empathy as an affective state isomorphic to the state of another person, encompassing a partial and experiential sharing of that person’s affect (Lamm et al., 2019; Hall & Schwartz, 2019 for overviews). Studies in recent years have already brought considerable advances in our understanding of the neural mechanisms underlying empathy (de Vignemont & Singer, 2006; Keysers & Gazzola, 2006; Lamm, Rütgen, & Wagner, 2019; Lockwood, 2016; Marsh, 2018; Preston & de Waal, 2002 for reviews; Jauniaux, Khatibi, Rainville, & Jackson, 2019; Lamm, Decety, & Singer, 2011 for meta-analyses). According to one influential account, the shared representations account, the experience of another individual’s emotion recruits neural processes that are (partially) functionally equivalent to those engaged during the first-hand experience of that emotion (Bastiaansen, Thioux, & Keysers, 2009; Lamm, Bukowski, & Silani, 2016; Lamm & Majdandžić, 2015 for reviews). Yet, apart from some general debate on the validity of this account (Zaki et al., 2016 for a review; but see also Zhou et al., 2020 for a recent preprint), there exists an explanatory gap regarding the relative contribution of somatosensory, compared to affective, brain regions to empathy.

Pain is widely used to study the neural underpinnings of empathy (Fan et al., 2011; Jauniaux et al., 2019; Lamm et al., 2011; Timmers et al., 2018 for meta-analyses). Classical first-hand pain processing is subdivided into two distinct brain networks, whose related brain activities map onto the first-hand experience of pain (Osborn & Derbyshire, 2010; Ploner, Gross, Timmermann, & Schnitzler, 2002; Jauniaux et al., 2019 for a meta-analysis; Tracey & Mantyh, 2007; Zaki, Wager, Singer, Keysers, & Gazzola, 2016 for reviews). Primary and secondary somatosensory cortices (S1/S2) encode information related to sensory-discriminative features of pain, such as location, timing or physical characteristics (Keysers, Kaas, & Gazzola, 2010; Vierck, Whitsel, Favorov, Brown, & Tommerdahl, 2013 for reviews). In turn, activity in anterior/midcingulate cortices (ACC/MCC) and anterior insula (AI) has been associated with affective-motivational aspects of pain, such as its subjective unpleasantness (Lockwood, 2016 for a review; Singer et al., 2004). While activation associated with the sensory-discriminative component is usually represented contralateral to the location of an applied stimulus (especially for S1, but also S2; Bingel et al., 2004; Haggard, Iannetti, & Longo, 2013; Ogino, Nemoto, & Goto, 2005; Omori et al., 2013; Ritter, Hebart, Wolbers, & Bingel, 2014), this has not been reported for the affective-motivational component (Lamm et al., 2011 for a meta-analysis). However, the relative importance of each component, and specifically the contribution of somatosensory processing to empathic pain experiences, remains controversial.

Numerous fMRI and EEG studies have demonstrated that receiving pain oneself and empathizing with another person in pain recruit overlapping activation in both of these pain processing components, providing possible evidence for shared representations (Lamm et al., 2011 for a meta-analysis; see Singer & Frith, 2005; Singer & Lamm, 2009 for reviews). For example, many studies continuously observed this overlap in bilateral AI and anterior MCC (aMCC), speaking for the affective-motivational component as the “core” of pain empathy processing (e.g. Benuzzi et al., 2018; Corradi-Dell’Acqua et al., 2011; Jackson et al., 2005; Singer et al., 2004; see Ding et al., 2019; Jauniaux et al., 2019 for meta-analyses). In addition, others reported overlapping activation in sensorimotor and somatosensory brain areas, highlighting the importance of the sensory-discriminative component for empathic pain experiences (e.g. Avenanti, Bueti, Galati, & Aglioti, 2005; Bufalari, Aprile, Avenanti, di Russo, & Aglioti, 2007; Gallo et al., 2018; Lamm, Nusbaum, Meltzoff, & Decety, 2007; Motoyama, Ogata, Hoka, & Tobimatsu, 2017; Riečanský & Lamm, 2019 for a review). Interestingly, results regarding the latter have only been reported when using specific types of paradigms.

To test the role of brain areas underpinning empathic responses more specifically and go beyond correlational evidence for shared activations, causal methods, such as psychopharmacological manipulations, have recently been used (Gallo et al., 2018). Placebo analgesia has been shown to reliably reduce first-hand pain using global (orally administered pill) or local (topically applied gel/cream) manipulations with no active pharmacological compound (Amanzio, Benedetti, Porro, Palermo, & Cauda, 2013 for a meta-analysis; Benedetti & Piedimonte, 2019; Colloca, Klinger, Flor, & Bingel, 2013; Wager & Atlas, 2015 for reviews; Corsi & Colloca, 2017). Rütgen, Seidel, Silani, et al. (2015) argued that if empathy for pain is indeed directly grounded in the experience of first-hand pain, placebo analgesia should also result in decreased empathy for pain. In three consecutive studies, they observed reduced self-reported empathy in participants in whom placebo analgesia had been induced (Rütgen et al., 2018; Rütgen, Seidel, Riečanský, et al., 2015; Rütgen, Seidel, Silani, et al., 2015). These results were later replicated by another group of researchers using the painkiller acetaminophen (Mischkowski et al., 2016). Imaging and EEG data further showed diminished activation during empathic pain processing in areas coding for the affective-motivational component (Rütgen, Seidel, Silani, et al., 2015) as well as reduced amplitudes of P2, an event-related potential (ERP) component (Rütgen et al., 2018; Rütgen, Seidel, Riečanský, et al., 2015). This component indexes neural computations related to the affective pain processing network and possibly also to somatosensory processing, as indicated by source localization studies (Cruccu et al., 2008; Perchet et al., 2012).

While these results suggest that empathy for pain is grounded in similar neural processes as first-hand pain (but see Lamm et al., 2019 and Zaki et al., 2016 for critical discussions), they also indicate that this neural sharing might only be partial and limited to a sharing of *affective* processes and representations. This brings back to the fore the unresolved issue about the role of the sensory-discriminative component in pain empathy (Fabi & Leuthold, 2017; Lamm et al., 2007; Loggia, Mogil, & Bushnell, 2008; Riečanský & Lamm, 2019 for a review; Singer et al., 2004). The previous studies from our lab did not report any variation in somatosensory activation by placebo analgesia, even when lowering statistical thresholds (Rütgen, Seidel, Riečanský, et al., 2015; Rütgen, Seidel, Silani, et al., 2015). This is surprising, given that placebo analgesia generally affects both components in first-hand pain (Benedetti, Mayberg, Wager, Stohler, & Zubieta, 2005; Wager & Atlas, 2015 for reviews). However, the experimental paradigm used in these studies may not have been tailored to provoke the engagement of somatosensory processes in the empathic experience, making their potential modulation by placebo induction difficult to discern (Keysers et al., 2010; Lamm et al., 2011). In fact, it has been suggested that *picture-based* empathy for pain paradigms directing the (visual and principal) attention of participants to the specific body part in pain, might be required to observe activation in somatosensory areas (e.g. visual input of a needle penetrating the hand; Timmers et al., 2018; Xiang, Wang, Gao, Zhang, & Cui, 2018 for overviews). Previous studies, however, employed a *cue-based* task, where facial expressions and abstract cues (Rütgen, Seidel, Silani, et al., 2015) or only abstract cues (Rütgen et al., 2018; Rütgen, Seidel, Riečanský, et al., 2015) indicated electrical stimulation given to the participants themselves or a second person. Thus, the task may not have been sufficiently sensitive to detect somatosensory modulation.

In this preregistered study, we therefore aimed to clarify the contribution of somatosensory processing in empathy for pain using an experimental paradigm allowing us to overcome the potential limitations of our previous research. To this end, we combined a causal experimental manipulation, consisting of a localized induction of placebo analgesia, with a paradigm putting a stronger emphasis on somatosensory aspects of the (empathic) pain experience than previous paradigms. More precisely, placebo analgesia was induced for one hand only, and participants’ attention was explicitly directed to the targeted hand by means of visual stimuli. In other words, we specifically optimized the study design in a way to maximize sensitivity for a potential placebo-driven modulation of somatosensory brain activity.

This motivated the following preregistered, directional hypotheses: First, we predicted reductions in first-hand and empathy for pain as well as unpleasantness ratings for the right hand, where placebo analgesia was induced, compared to the left hand acting as a control. Second, we hypothesized that the sensory-discriminative component of pain empathy would be modulated in a similar fashion by placebo analgesia as the affective-motivational component – i.e., that neural responses related to the right hand would be reduced in S1 and S2 compared to the left hand – and that this would trigger correspondingly reduced neural responses in bilateral AI and aMCC.

## 2 Materials and methods

### 2.1 Data and code availability statement

The data was newly acquired for the present study. Unthresholded statistical maps will be made available via an online repository upon acceptance and stimuli templates for the pain task are uploaded within the Open Science Framework (OSF) project (osf.io/2q3zu/).

### 2.2 Preregistration

We report how we determined our sample size, all data exclusions, all manipulations, and all measures in the study. This study was preregistered on the OSF prior to any creation of data (Hartmann, Rütgen, Sladky, & Lamm, 2018; preregistration: osf.io/uwzb5; addendum: osf.io/h7v9p) and was designed to extend and specify the results of Rütgen, Seidel, Silani, et al. (2015) in regard to somatosensory sharing. Methods reported below are therefore reproduced partly verbatim from the preregistration. Note that the preregistered plan contains a second research question that is not part of the present paper but will be reported elsewhere. In the following methods and results, we clearly distinguish preregistered procedures and analyses from those added post hoc.

### 2.3 Participants

Participants were recruited by means of flyers and online advertising in Vienna, Austria and via an existing database of study participants. Upon interest, they were screened by means of an online questionnaire (see A.1 in Supplement A for detailed information regarding exclusions). An *a priori* power analysis using G*Power 3 (Faul et al., 2007) was conducted using a conservative average of the lowest effect sizes from previous placebo empathy analgesia studies (one-tailed paired *t*-test, Cohen’s *d* = .79 to .44 for self-report and .40 to .39 for affective brain areas; Rütgen, Seidel, Riecansky, & Lamm, 2015; Rütgen, Seidel, Silani, et al., 2015) to calculate the needed sample size to detect a medium effect size of *d =* .40 at a standard error probability of α = .05 with a power of 1-ß = 0.8. This yielded a sample size of 41 participants. However, considering that the modulation of placebo analgesia might not be equal for somatosensory compared to previously reported affective brain regions, a total of 45 placebo responders was set as the stopping-rule. The exclusion of nonresponders in regard to the placebo manipulation was crucial to obtain a sample of participants showing a robust localized, first-hand placebo analgesia effect, in order to investigate a transfer of this effect to empathy. We originally included three measures to identify nonresponders in our preregistration, as per the criteria in Rütgen, Seidel, Silani, et al. (2015). During data collection, we uploaded an addendum to include a fourth measure we had previously overlooked. This measure was not possible in the previous study we oriented our procedures on but was added due to our within-subjects design in order to better identify nonresponders, maximize the placebo responsiveness of the final sample and bolster the interpretability of our results. We had not observed or analysed any of the collected data when preregistering this addendum. Importantly, this was also the criterion that identified almost all of the nonresponders (see also A.2 in Supplement A).

Our final sample included 22 males and 23 females (Age: *M* ± *SD* = 23.84 ± 2.73 years, range = 19-32; all right-handed with laterality quotients (LQs) ≥ 80 and normal or corrected-to-normal vision). We purposefully recruited only strongly right-handed participants and did not counterbalance the location where placebo analgesia was induced between participants to avoid laterality problems in our fMRI analyses, as well as to increase sample homogeneity and comparability of the induction procedure. Before the commencement of the study, five pilot participants were tested to confirm the existence of a localized, first-hand placebo analgesia effect and improve study procedures, but these datasets were not included in the final sample. All participants gave written consent at the outset of each session. The study was approved by the ethics committee of the Medical University of Vienna (EK-Nr. 661/2011) and performed in line with the latest revision of the Declaration of Helsinki (2013).

### 2.4 Procedure

The study consisted of two parts: First, participants came alone for a one-hour session to the lab, where they filled out questionnaires on a computer, and had photos of their hands taken that were used as individualized stimulus material for the scanning session. After an average interval of 32.86 ± 29.16 (*M* ± *SD*) days, participants came to the MRI scanner where they took part in the main experiment. Each one arrived together with a second person (who was a female confederate of similar age invited by the experimenters acting as a second participant, as per Rütgen, Seidel, Silani, et al., 2015). The experimenter explained to both that the goal of the study was to investigate brain activity associated with a local anesthetic in the form of a medical gel. Furthermore, it was made clear that only one person, i.e. the participant, would receive this medication on the right hand and complete the tasks inside the scanner, while the confederate would not receive any medication and complete the same tasks on a computer next to the scanner.

After signing the consent form and the MR-safety questionnaire, the confederate was asked to wait outside the control room while an individual psychophysical pain calibration was performed with the participant. This was done to determine the maximum level of tolerable pain and to specify average subjective values for very painful (rating of 7 on a scale from 0 = not painful to 8 = extremely painful), medium painful (rating of 4) and not painful, but perceivable (rating of 1) stimulation. As pain tolerances can vary depending on the body part and handedness (Murray & Safferstone, 1970; Pud et al., 2009), we calibrated each hand individually to match the stimulation intensities and subjective pain levels for each hand. The hand calibrated first was counterbalanced across participants. To this end, an electrode was attached to the dorsum of each hand using medical tape. Electrical stimulation of various strengths (stimulus duration = 500 ms) was administered using the procedure employed by Rütgen, Seidel, Silani, et al. (2015), with two rounds going from very low (0.05 mA) to continuously higher stimulation until the participant indicated the last received stimulus as an ‘8’, after which each round was terminated. This was followed by a third round of stimuli with random intensity in the before calibrated range. Short breaks between the stimuli and longer breaks of a few minutes between the rounds ensured an independent rating of each stimulus unbiased by previous one(s). Participants were instructed to rate each stimulus as intuitively but also as accurately as possible. Input intensities for the task were the individual average ratings for painful (rating of 7) and non-painful (rating of 1) stimulation given during calibration, separately for the left and right hand. Those were 0.64 ± 0.67 (*M* ± *SD*) mA (left hand) and 0.53 ± 0.36 mA (right hand) for painful, and 0.09 ± 0.06 mA (left hand) and 0.10 ± 0.06 mA (right hand) for non-painful sensations. We compared values for painful and non-painful stimulation separately for left and right hands using two paired *t*-tests in order to investigate differences in pain tolerance between the hands (analysis not preregistered; Murray & Safferstone, 1970; Pud et al., 2009). Stimulation intensities did not differ between the two hands (pain: *t*(44) = 1.59, *p* = .117; no pain: *t*(44) = -0.99, *p* = .325). In general, electrical stimulation was delivered using a Digitimer DS5 Isolated Bipolar Constant Current Stimulator (Digitimer Ltd, Clinical & Biomedical Research Instruments).

Next, a medical student in a white lab coat posing as the study doctor introduced the medication as a “powerful local anesthetic” and gave information on its effects and possible side effects. Participants were told that the medication would be effective after a 15-20 minutes waiting period and then remain stable for 2-3 hours. The study doctor attached a white paper bracelet to the right wrist of the participants (as a visual reminder which hand received the “medical treatment”), then applied and rubbed in the placebo gel on the dorsum of the right hand. On the left hand, participants were told that a control gel with no active ingredients was applied. In reality, both gels contained nearly the same basic ingredients of a standard skin gel with no active pharmacological components (see A.3 in Supplement A for exact ingredients). In matching the two gels, we aimed for clearly recognizable visual and olfactory distinction, but the same tactile feeling and hydrating properties, and adhered to previously used procedures inducing placebo analgesia with topical creams and gels (e.g. Benedetti et al., 1999; Bingel et al., 2006; Geuter et al., 2013; Schenk et al., 2014; Tinnermann et al., 2017). After the application, the participant was led outside the control room to (ostensibly) wait for the “medication” to take effect and was told that the confederate would undergo the same pain calibration in the meantime. During the waiting period, the participant was instructed regarding the pain task. After 15 minutes, the participant returned to the control room and was told that the effectiveness of the medication would now be verified using a “pain test”. Here, we employed a classic conditioning procedure to amplify the effects of the placebo. After removal of excess gel and disinfection with 70% isopropyl rubbing alcohol, one electrode was again attached to the dorsum of each hand, using the same placement as during calibration. Participants were told that they would be getting stimulation on both hands that they had judged as “painful” before, and were asked to rate how painful it felt for them. On the left (control) hand, participants indeed received stimulation with a prior subjective rating of 7 (“very painful”), on the right (placebo) hand, however, they covertly received stimulation with a prior rating of 4 (“medium painful”) to suggest substantial pain relief by the medical gel. All participants completed at least two conditioning rounds (in the first round, three successive stimuli were given, in subsequent rounds four), and were given oral feedback after each round by the experimenter, namely that their ratings on the left/control hand were similar to their ratings during calibration, but the ratings on the right/placebo hand had decreased substantially. If participants rated the stimuli on the right/placebo hand greater than 5 and/or the stimuli on the left/control hand lower than 5, the conditioning round was deemed unsuccessful and repeated up to a maximum of four times. After unsuccessful rounds, stimuli were slightly adjusted for the next round(s) to increase the contrast between the two hands, i.e. increasing intensities for the left hand and/or decreasing intensities for the right hand. This was done without knowledge of the participants, who thought they received the same level of stimulation on both hands at all times.

Afterwards, the participant and the confederate were led into the scanner room and the confederate was seated on a table with a computer screen, keyboard and headphones next to the scanner. Following general adjustments, the participant completed two runs (22 minutes each) of the pain task, and one run of another task (not reported here) in a fixed order. The rating hand of the participants was counterbalanced between but kept constant within participants over all tasks. Upon completion of all tasks, the experimenter went inside the scanner room pretending to get the confederate, after which the field map and structural image were acquired. After scanning, participants filled out post-experimental questionnaires. They received a compensation of 50 Euros for taking part in the whole study and an aliquot amount if they dropped out earlier. The overall scanning session took ∼ 4 hours, of which participants spent around 80 minutes lying in the scanner.

### 2.5 Pain task

To induce pain, we used short-lasting painful and non-painful electrical stimulation delivered to the right and left hands of the participant or confederate in different trials. By adding a non-painful control stimulation, our effects can be more specifically attributed to pain processing. Domain-general aspects (such as generalized perceptual or behavioral responses, including stimulus-directed attention) related to stimulus presentation are explicitly eliminated by this approach (Petrovic, Kalso, Petersson, & Ingvar, 2002; Rütgen, Seidel, Silani, et al., 2015). The pain task was implemented in MATLAB R2017b (Mathworks) using the Cogent 2000 Toolbox Version 1.33 (http://www.vislab.ucl.ac.uk/cogent_2000.php). Participants saw either pictures of their own hands (with the right/placebo hand wearing a white bracelet) or the confederate’s hands from an egocentric perspective on black background, depending on who would receive the next stimulation (see Figure 1 and A.4 in Supplement A).

**Figure 1.**
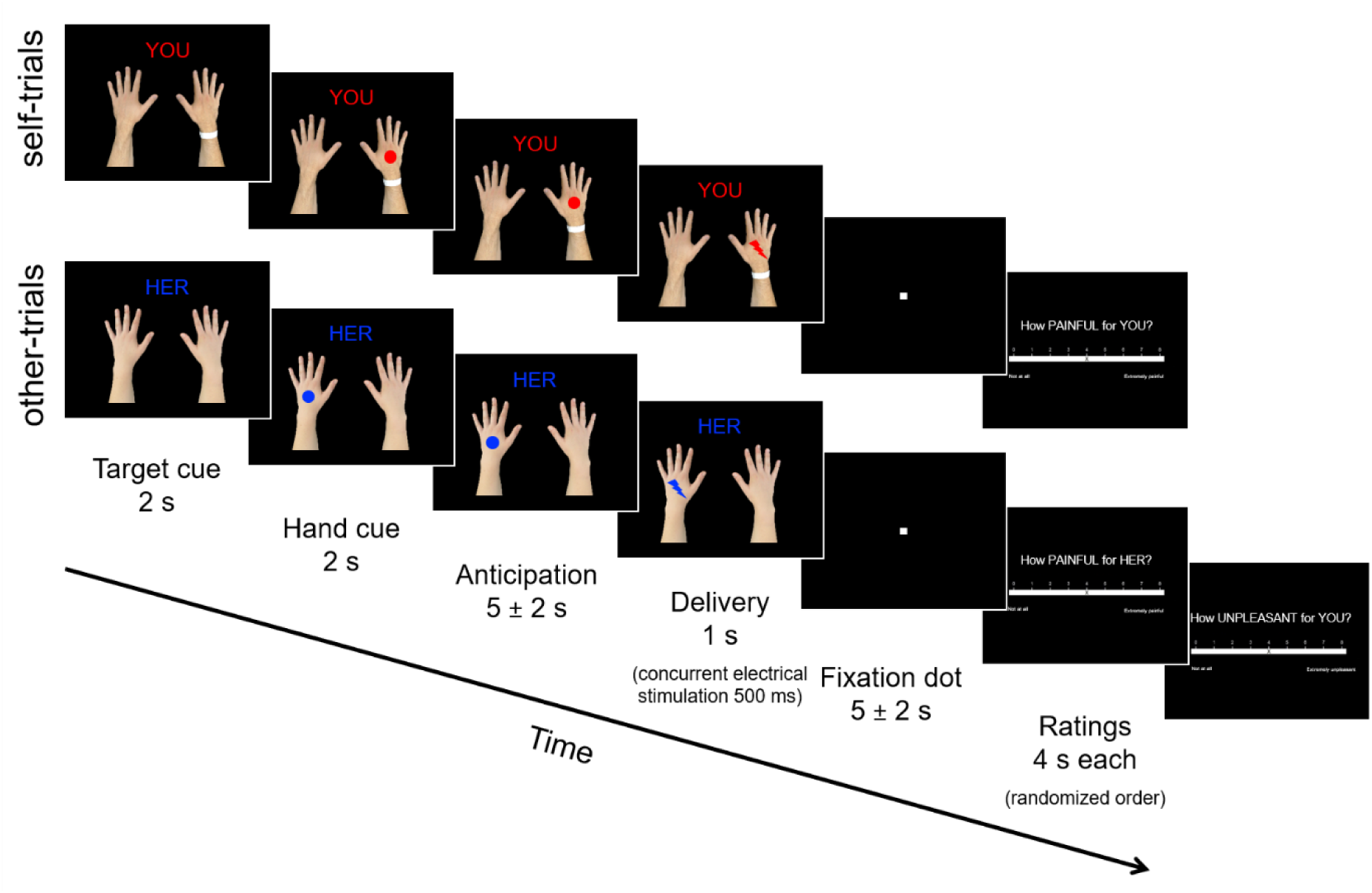
Overview of the pain task. As part of a 2×2×2 full-factorial design, participants either received painful (red icons) or non-painful (blue icons) electrical stimulation themselves (seeing their own hands; self-trials) or witnessed a second person receiving such stimulation (seeing the confederate’s hands; other-trials). Prior to the task, all participants had undergone a localized placebo analgesia induction on their right hand, while the left hand acted as each participant’s individual control. In half of all trials, subjective ratings were collected after stimulus delivery for self- and other-related pain intensity, as well as self-related unpleasantness when observing the other person in pain.

Each trial began with the written German words “DU” (“YOU”, self-trials) or “SIE” (“HER”, other-trials) in either red or blue (for painful or non-painful stimulation, respectively), indicating the target and the intensity of the next stimulation (target cue; 2000 ms). Then, a circle icon in the same color of the word was shown on the hand receiving the next stimulation (hand cue; 2000 ms). After a jittered anticipation period (5000 ± 2000 ms, evenly distributed in 500 ms steps) simultaneously displaying the hands with the two cues, the circle changed into a lightning icon of the same color, indicating stimulus delivery (duration of electrical stimulus = 500 ms, display of delivery cue = 1000 ms). This was followed by a jittered waiting period (5000 ± 2000 ms, evenly distributed in 500 ms steps), depicting a white dot on black background. In half of all trials, stimulation delivery was followed by a rating period (4000 ms per question; appearance of the rating phase was determined by four pseudorandomized sequences previously created). During self-trials, participants were asked how painful the stimulus had felt for them. During other-trials, participants were asked two questions tapping into different aspects of empathy (Coll et al., 2017; Lamm & Majdandžić, 2015), namely (1) how painful the stimulus was for the other person (cognitive-evaluative aspect) and (2) how unpleasant it was for the participant him- or herself to witness the other person receiving such stimulation (affective-sharing aspect). The two empathy questions always appeared in a random order. Questions were rated on visual analogue scales from 0 = “not perceivable at all” to 8 = “extremely painful/unpleasant”. A 2000 ms inter-trial-interval screen depicting a white dot on black background was shown before the start of the next trial. Participants completed 128 trials with an average duration of 21/25 s (self-trials/other-trials) per trial, 64 trials per run and 16 trials per condition, with trials appearing in one out of four pseudorandom orders previously created.

### 2.6 MRI data acquisition

MRI data was acquired using a 3 Tesla Siemens Magnetom Skyra MRI-system (Siemens Medical, Erlangen, Germany), equipped with a 32-channel head coil. The functional scanning sequence included the following parameters: Echo time (TE)/repetition time (TR) = 34/1200 ms, multi-band acceleration factor = 4, flip angle = 66°, interleaved multi-slice mode, interleaved acquisition, field of view = 210 mm, matrix size = 96×96, voxel size = 2.2×2.2×2.0 mm^3^, 52 axial slices of the whole brain coplanar the connecting line between anterior and posterior commissure, and slice thickness = 2 mm. Functional volumes were acquired in two runs (and one run for another task), with small breaks in between the three runs. Structural images were acquired using a magnetization-prepared rapid gradient-echo sequence (TE/TR = 2.43/2300 ms, ascending acquisition, field of view = 240 mm, single shot multi-slice mode, 208 sagittal slices, voxel size = 0.8×0.8×0.8 mm^3^, flip angle = 8°, slice thickness = 0.8 mm).

### 2.7 Behavioral data analysis

The analysis workflow of the behavioral and fMRI data is summarized in Figure 2 and referred to throughout the following methods and results. Statistical analyses were performed in RStudio Version 3.6.1 (R Core Team, 2019; for analysis and plotting functions see A.5 in Supplement A). We conducted all our preregistered *t*-tests one-tailed due to *a priori* directional hypotheses.

**Figure 2.**
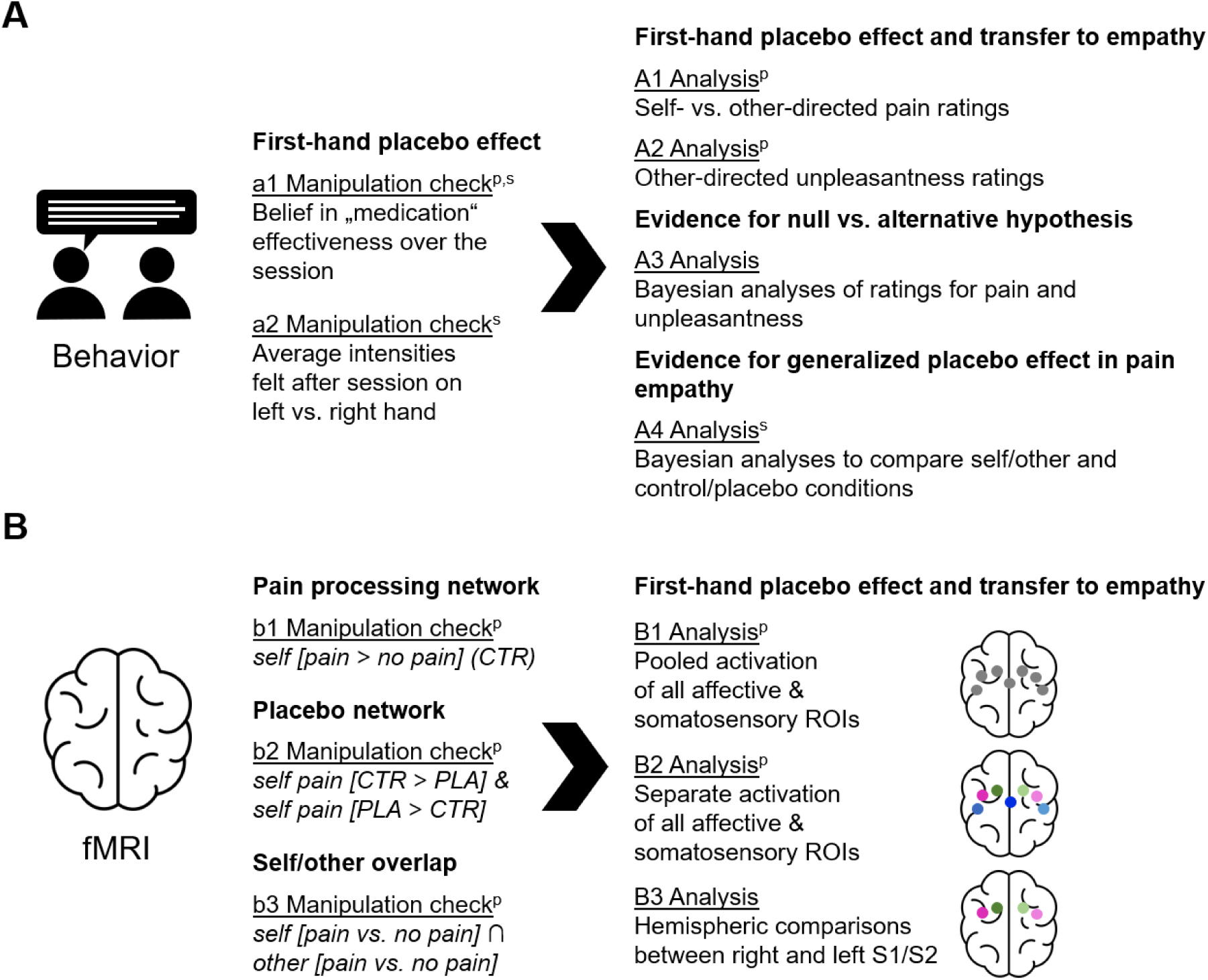
Overview of the analysis workflow. A) For the behavioral data, we explored the validity of our design using two manipulation checks (a1+a2; reported in A.6 in Supplement A). Then, we conducted four analyses to evaluate the evidence for a first-hand localized placebo analgesia effect and a transfer of this effect to empathy using the ratings collected from the task (A1-A4; A4 is reported in Supplement B). B) Regarding the fMRI data, we used three manipulation checks to establish the validity of our pain task (b1), the typical placebo analgesia network (b2) and the previously reported self-other overlap in brain activity related to first-hand and empathy for pain (b3). For our main analyses, we employed a region of interest (ROI) approach to evaluate the evidence for a first-hand localized placebo analgesia effect and a transfer of this effect to empathy in seven ROIs: anterior midcingulate cortex, bilateral anterior insula, as well as bilateral primary (S1) and secondary somatosensory cortex (S2). This was first done using pooled activation of all ROIs (B1) and then analyzing each ROI separately (B2). Finally, we gathered further evidence for a first-hand localized placebo effect and absence of a transfer to empathy using a hemispheric comparison analysis (B3). Preregistered analyses are marked with ^p^, analyses in the supplement are marked with ^s^; PLA = placebo; CTR = control.

#### 2.7.1 Preregistered analyses

We implemented a within-subjects, full-factorial design with three factors of two levels each (*treatment*: placebo vs. control hand, *target*: self vs. other, *intensity*: pain vs. no pain). Two parametric repeated-measures analyses of variance (ANOVAs) were used to analyze the results. In the first ANOVA (analysis A1 in Figure 2), the dependent variable was the self- and other-related pain ratings. A second ANOVA (analysis A2 in Figure 2) included the unpleasantness ratings as the dependent variable (omitting the factor *target*, as unpleasantness ratings were only collected in the empathy condition). For each ANOVA, we then computed planned comparisons using paired *t*-tests.

#### 2.7.2 Post hoc analyses

Due to the unexpected “null” finding of no transfer of the first-hand placebo effect to empathy, we aimed to gather further relative evidence for the null vs. the alternative hypothesis, using a Bayesian approach (e.g. Wagenmakers et al., 2018). This was realized with three Bayesian paired *t*-tests (analysis A3 in Figure 2) mirroring the above preregistered analyses. We used a standard Cauchy (0,1) prior as the effect size (indicating a 50% chance to observe an effect size between -1 and 1; e.g. Rouder et al., 2009). Note that Bayesian *t*-tests produce a Bayes Factor comparing the relative evidence between the alternative and null hypothesis (BF_10_, H_1_ vs. H_0_; Giolla & Ly, 2019). In interpreting these values, a BF_10_ < 3 has been suggested to indicate weak evidence, a BF_10_ > 3 positive evidence, and BF_10_ > 150 very strong evidence for the alternative hypothesis (Jarosz & Wiley, 2014). Evidence for the null compared to the alternative hypothesis (BF_01_, H_0_ vs. H_1_) was computed as BF_01_ = 1/BF_10_. For an additional analysis exploring the existence of any placebo-related downregulatory effect (analysis A4 in Figure 2) as well as results and discussion, see Supplement B.

### 2.8 fMRI data preprocessing and analysis

#### 2.8.1 Preprocessing and first-level analysis

To preprocess and statistically analyze the fMRI data, the software Statistical Parametric Mapping (SPM12, https://www.fil.ion.ucl.ac.uk/spm/software/spm12/, Wellcome Trust Centre for Neuroimaging) running on MATLAB Version R2015b (Mathworks) was used. All brain regions were labelled with the SPM Anatomy toolbox version 2.15 (Eickhoff et al., 2005). Preprocessing of the functional volumes included slice timing (reference = middle slice; Sladky et al., 2011), realignment with the participant-specific fieldmap, coregistration of structural and functional images, segmentation into gray matter, white matter (WM) and cerebrospinal fluid (CSF) tissues, spatial normalization, and spatial smoothing by convolution with an 8 mm full-width at half-maximum (FWHM) Gaussian Kernel. The first-level design matrix of each participant contained eight regressors for anticipation (combining target + hand cues), eight for delivery and one for all rating phases, leading to 17 regressors. The different conditions were modeled in an event-related fashion and convolved with SPM12’s standard canonical hemodynamic response function. Additional nuisance regressors included six realignment parameters and two regressors modeling WM and CSF for each of the runs (the latter two were extracted using the REX toolbox; Duff, Cunnington, & Egan, 2007). We excluded 1-2 trials in four participants from analysis post hoc due to technical malfunctioning of the pain stimulator (e.g. missing stimulation in a pain trial).

#### 2.8.2 Manipulation checks

We preregistered three manipulation checks testing (i) the validity of our design, (ii) the success of the placebo analgesia induction and (iii) the existence of overlapping activation for first-hand and empathy for pain (manipulation checks b1-b3 in Figure 2). To this end, eight contrast images were created for each participant (these were not specified in the preregistration, but we adhered to the procedure used in our previous study, modeling the whole time phase from the first cue and anticipation phase until one second after delivery onset; Rütgen, Seidel, Silani, et al., 2015). We then calculated a full factorial model within a flexible factorial framework in SPM12 using the within-subjects factors *treatment* (placebo vs. control hand) and *condition* (combining the factors *target* (self, other) and *intensity* (pain, no pain)), as well as the between-subjects factor *subject*. We determined significance using cluster-level inference. To correct for multiple comparisons, we calculated the cluster extent threshold by means of “CorrClusTh.m”, an SPM extension script (Thomas Nichols, University of Warwick, United Kingdom & Marko Wilke, University of Tübingen, Germany; http://www2.warwick.ac.uk/fac/sci/statistics/staff/academicresearch/nichols/scripts/spm/).

First, we used the contrast *[self - pain > self - no pain]* of the control hand to evaluate whether our design robustly activated brain areas associated with pain processing as in previous studies (manipulation check b1 in Figure 2). This check is reported at a cluster probability of *p* < .05 (familywise-error (FWE)-corrected cluster-forming threshold of *k* = 188, initial cluster-defining threshold *p* < .001 uncorrected).

Second, we aimed at showing that the placebo analgesia induction activated a widespread network previously identified in placebo analgesia studies (manipulation check b2 in Figure 2; see e.g. Atlas & Wager, 2012 for a summary). Here we used small volume correction (SVC), with a threshold of *p* < .05 FWE-corrected at peak-level, on the contrasts *[self - placebo hand > self - control hand]* and *[self - control hand > self - placebo hand]*, using only the pain conditions (initial threshold: *p* < .001 uncorrected). This approach was directly motivated by previous studies (e.g. Bingel et al., 2007; Eippert et al., 2009; Geuter et al., 2013; Wager et al., 2011; Zubieta et al., 2005) and chosen to maximize sensitivity of the analyses. In accordance with these studies, we analysed spheres around MNI coordinates used in the study by Rütgen, Seidel, Silani, et al. (2015), as this study’s overall design closely matched the present one, and as this allowed us to compare data from within the lab ([±x, y, z]; size of sphere): Dorso-lateral prefrontal cortex (DLPFC; [±36, 13, 39]; 15 mm), S2 ([±39, -15, 18]; 10 mm), insula (anterior [±33, 18, 6] and posterior [±44, -15, 4] part; both 10 mm), dorsal (dACC; [±3, 6, 36]; 10 mm) and rostral ACC (rACC; pregenual [±10, 32, -8] and subgenual [±6, 30, -9] parts; both 10 mm), ventral striatum ([±9, 6, -3]; 6 mm), thalamus ([±12, -18, 3]; 6 mm), and periaqueductal gray ([0, -32, -10]; 6 mm).

Thirdly, to check that the design evoked empathic responses that overlapped with the first-hand experience of pain, we performed a conjunction analysis between self- and other-related conditions using the contrast *[self - pain > self - no pain]* ∩ *[other - pain > other - no pain]* for the control hand only (manipulation check b3 in Figure 2). However, this preregistered check did not reveal any significant clusters, when using a whole brain and FWE-cluster-corrected approach. The following checks were therefore added post hoc: To investigate this overlap in previously reported affective brain regions related to empathy, we adopted a SVC approach on three ROIs from an independent meta-analysis using the above two contrasts (Lamm et al., 2011): left AI [-40, 22, 0], right AI [39, 23, -4] and aMCC [-2, 23, 40] (all 10 mm), again *p* < .05 FWE-corrected at peak-level. Furthermore, to maximize sensitivity and detect any overlap between self- and empathy-related conditions, we additionally reported the same conjunction with contrasts averaging over both hands.

#### 2.8.3 Preregistered analyses

To test our hypothesis of a somatosensory-specific transfer of placebo analgesia to empathy for pain, we conducted ROI analyses in bilateral AI and aMCC (see coordinates above taken from Lamm et al., 2011), as well as in bilateral S1 and S2 (left S1: [-39, -30, 51]; right S1: [36, -36, 48]; left S2: [-39, -15, 18]; right S2 [39, -15, 18]; see Figure 3). S1/S2 coordinates were taken from independent findings investigating first-hand somatosensory pain perception (Bingel et al., 2004 for S1, 2007 for S2). We created 10 mm spheres around each coordinate with MarsBaR (Brett et al., 2002) and then extracted parameter estimates for each ROI from the first-level contrast images of each participant and for each condition using REX (Duff et al., 2007). The specific coordinates and sphere sizes for the ROI analyses were not preregistered, but we again adhered to procedures used in Rütgen, Seidel, Silani, et al. (2015). ROI analyses were conducted in RStudio Version 3.6.1 (R Core Team, 2019). Mimicking the behavioral analysis, we implemented the same within-subjects, full-factorial design with three factors (*treatment, target, intensity*) of two levels each, and the additional factor *ROI* with seven levels (pooled activation of lAI, rAI, aMCC, lS1, rS1, lS2, and rS2; analysis B1 in Figure 2). In the pooled ANOVA, Mauchly’s test for sphericity was significant for the main effect of ROI and all interactions with the factor ROI, which is why those results are reported using Greenhouse Geisser sphericity correction. Due to the significant main effect of ROI and interactions with the factor ROI in the initial four-way ANOVA, we proceeded with our preregistered analysis plan by computing separate ANOVAs and planned comparisons for each of the ROIs (analysis B2 in Figure 2). Again, all preregistered *t*-tests were conducted one-tailed.

**Figure 3.**
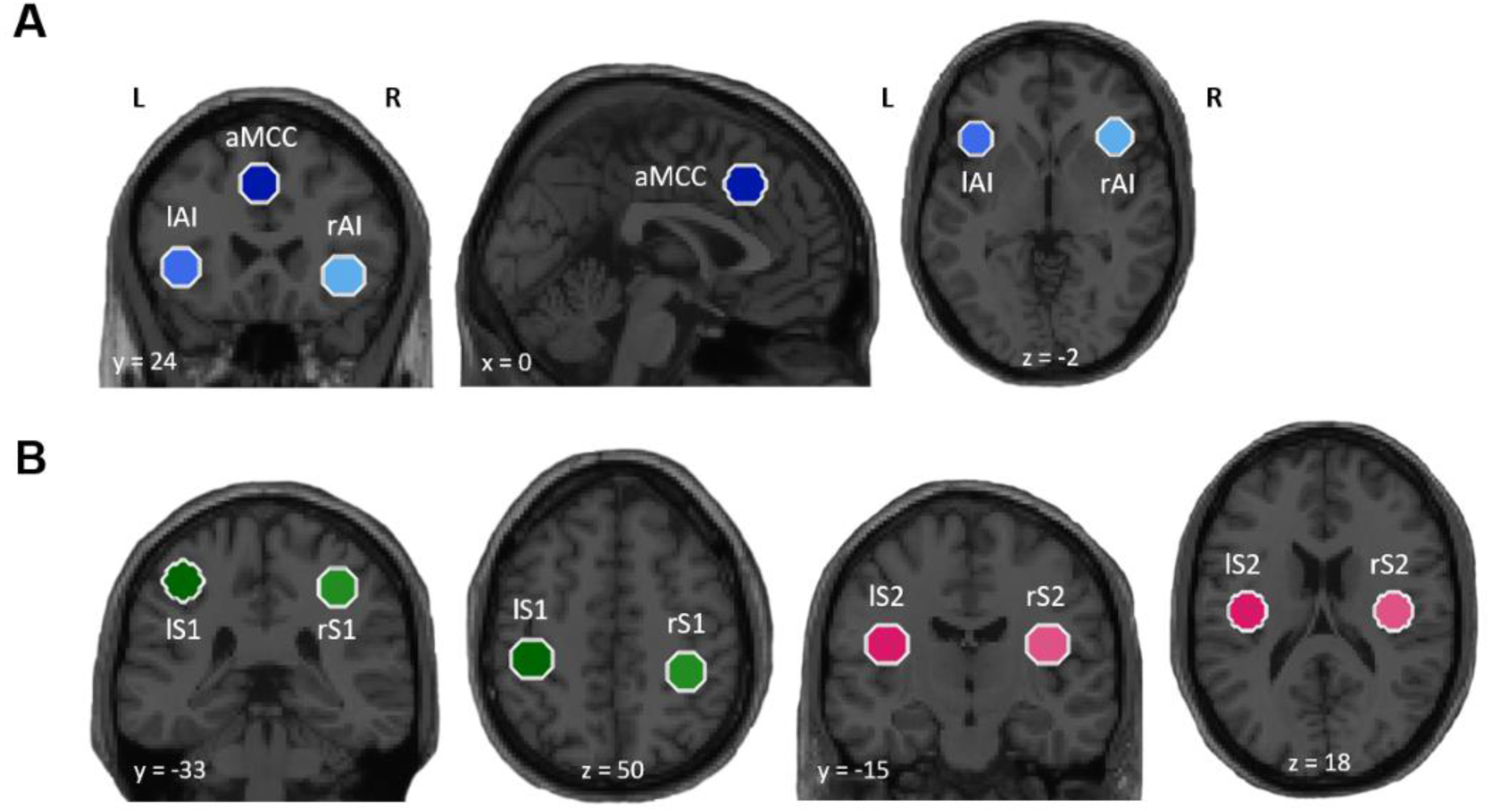
Overview of the seven regions of interest (ROIs) used in the main analysis. We analysed the transfer of the first-hand placebo effect to empathy for pain in A) three affective and B) four somatosensory brain regions (all 10 mm spheres; MNI coordinates [x, y, z]: left/right anterior insula (lAI: [-40, 22, 0]; rAI: [39, 23, -4]), anterior midcingulate cortex (aMCC: [-2, 23, 40]), left/right primary somatosensory cortex (lS1: [-39, -30, 51]; rS1: [36, -36, 48]; left/right secondary somatosensory cortex (lS2: [-39, -15, 18]; rS2: [39, -15, 18]; anatomical brain regions were confirmed with the SPM Anatomy toolbox version 2.15 by Eickhoff et al., 2005). Bilateral AI and aMCC coordinates were taken from an independent meta-analysis on networks involved in (empathic) pain (Lamm et al., 2011), while bilateral S1/S2 coordinates were taken from two studies investigating somatosensory pain perception (Bingel et al., 2004 for S1, 2007 for S2). L = left hemisphere, R = right hemisphere.

#### 2.8.4 Post hoc analyses

Our preregistered main analysis tested for the difference between placebo and control hand in each ROI, e.g. activation differences in right S1 during stimulation of left (contralateral) control hand vs. right (ipsilateral) placebo hand. However, although stimulation of one body site often evokes bilateral activation, most studies investigating somatosensation of noxious and non-noxious stimuli report a strong contralateral bias, i.e. a location coding in the contralateral hemisphere for S1 and S2 (Bingel et al., 2003; Bingel et al., 2004; Ogino et al., 2005; Tamè et al., 2012; Wager et al., 2004). Thus, our preregistered analysis approach was not optimized to deal with possible laterality issues in these two regions. Therefore, to gather additional evidence that our participants had in fact a first-hand placebo analgesia effect that was localized, or in other words, specific for the right hand (and to ensure that this effect did in fact not transfer to empathy), we conducted a hemispheric comparison (analysis B3 in Figure 2) aimed at directly contrasting brain activation in the corresponding contralateral hemispheres related to each hand (e.g. activation in right S1 during stimulation of left control hand vs. activation in left S1 during stimulation of the right placebo hand). Mirroring previous approaches, we used the pain conditions only (e.g. Eippert et al., 2009; Rütgen, Seidel, Silani, et al., 2015; Zubieta et al., 2005). We focused this analysis on S1 and S2, since both have been found to provide spatial information of painful and non-painful stimulation in the hemisphere contralateral to the stimulated body side (Bingel et al., 2003). However, while S1 is more often reported in relation to general stimulation (Keysers et al., 2004; Ploner et al., 2000), S2 additionally seems to encode stimulus intensity and play a greater role in the processing of pain (Lockwood et al., 2013). Furthermore, involvement of S2 in placebo analgesia mechanisms has been reported, making S2 an especially optimal candidate for testing the localized first-hand placebo analgesia effect in our study (Bingel et al., 2003, 2007; Bingel et al., 2004; Eippert et al., 2009; Geuter et al., 2013; Price, Craggs, Nicholas Verne, Perlstein, & Robinson, 2007; Schenk et al., 2014; Wager et al., 2011, 2004). Our aim for this analysis was thus to directly compare the two hemispheres of both regions with each other, but only considering activation related to each contralateral hand. To this end, we used the previously extracted parameter estimates of left and right S1 and S2. For each region and hemisphere, we subtracted activation related to the ipsilateral hand from activation related to the contralateral hand (e.g. right S1 = activation related to left control hand stimulation - activation related to right placebo hand; left S1 = activation related to right placebo hand - activation related to left control hand). Then, we used these subtracted values to compare activation in left S1 related only to the right placebo hand with activation in right S1 related only to the left control hand (and the same for S2) via two-tailed paired *t*-tests. This was done separately for self- and other-related stimulation.

## 3 Results

### 3.1 Behavioral results

The two manipulation checks showed a strong belief in the effectiveness of the placebo gel over the course of the session and a robust behavioral placebo effect even afterwards (manipulation checks a1 and a2 in Figure 2; see A.6 and Figure A1 in Supplement A).

#### 3.1.1 Preregistered analyses

To evaluate the existence of a localized first-hand placebo analgesia effect for pain as well as the transfer of this effect to other-related pain and self-experienced unpleasantness, we calculated two repeated-measures ANOVAs. The first ANOVA using self- and other-related pain ratings revealed all main effects and interactions to be significant (analysis A1 in Figure 2; see Table A1 in Supplement A and Figure 4 for an overview of all behavioral ratings). Planned comparisons showed a significant placebo analgesia response for self-related but not for other-related stimulation (self: *t*(44) = 9.49, *p* < .001 one-tailed, *M*_*diff*_ = 1.619, 95% CI_meandiff_ [1.28, 1.96], Cohen’s *d*_*z*_ = 1.42; other: *t*(44) = -0.17, *p* = .435 one-tailed, *M*_*diff*_ = 0.018, 95% CI_meandiff_ [-0.23, 0.19], Cohen’s *d*_*z*_ = 0.03). Indeed, the mean ratings were decreased in the placebo compared to the control hand for first-hand stimulation (see Figure 4). This was not the case for pain empathy, where the mean ratings for the two hands were similar. The magnitude of the placebo effect, i.e. the difference between placebo and control hand, was significantly higher in the self, compared to the other (self vs. other: *t*(44) = -8.22, *p* < .001 one-tailed, *M*_*diff*_ = -1.64, 95% CI_meandiff_ [-2.04, -1.24], Cohen’s *d*_*z*_ = 1.23).

**Figure 4.**
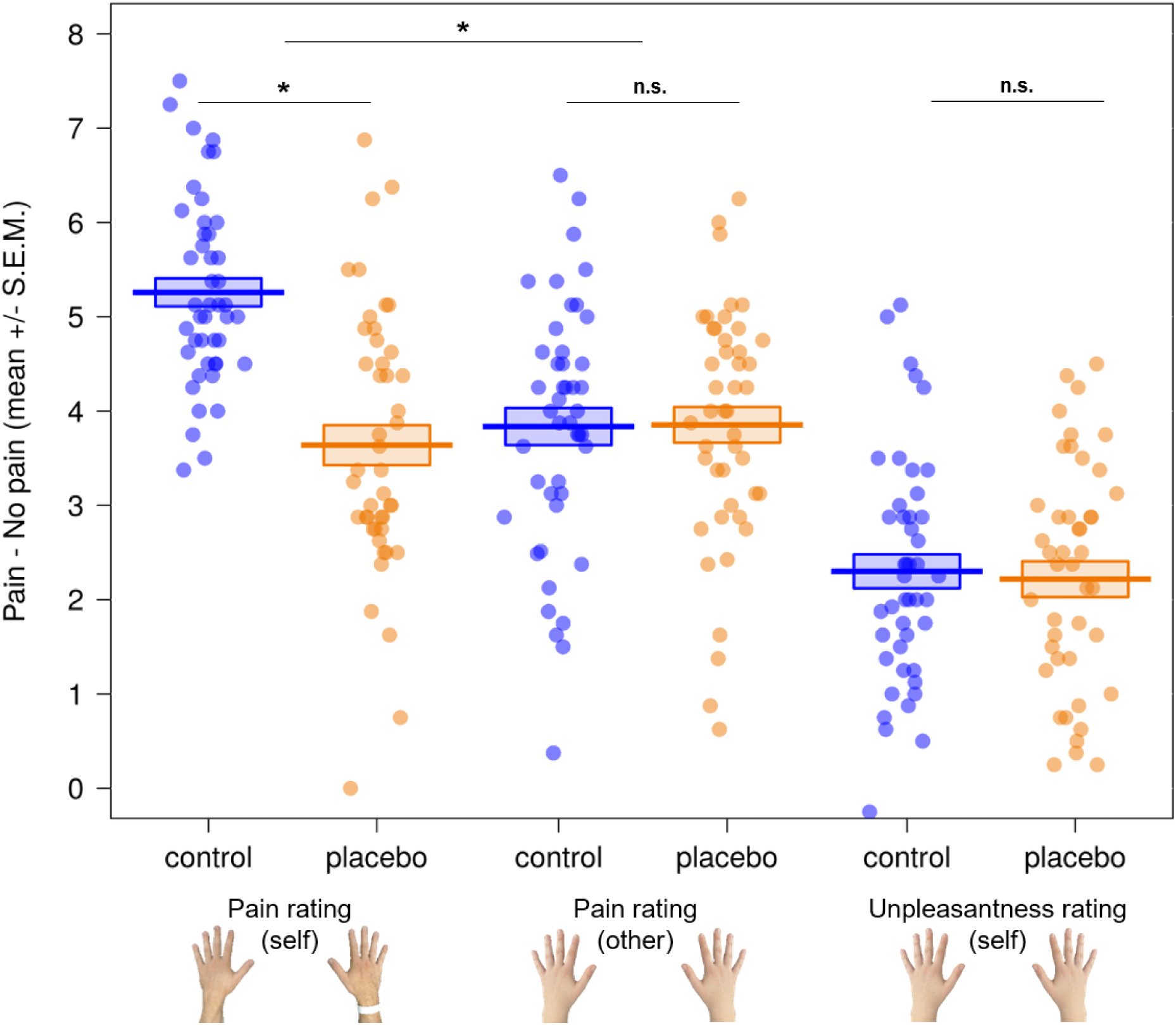
Behavioral results of the pain task. Participants rated electrical stimulation they either received themselves or witnessed another person receiving (displayed here as an index of the ratings for pain - no pain conditions). Using paired *t*-tests, we observed a significant placebo effect for self-related pain ratings, but no transfer to other-related pain or self-experienced unpleasantness ratings when observing the other in pain. * *p* < .05; n.s. = not significant; S.E.M. = standard error of the mean.

The second ANOVA, using ratings of the participants’ own unpleasantness while watching the confederate receiving stimulation, showed similar results, with a main effect of intensity but no hand x intensity interaction (analysis A2 in Figure 2; see Table A2 in Supplement A). The planned comparison indicated no placebo analgesia effect related to one’s own unpleasantness (*t*(44) = 0.69, *p* = .245 one-tailed, *M*_*diff*_ = 0.084, 95% CI_meandiff_ [-0.16, 0.33], Cohen’s *d*_*z*_ = 0.10). Participants experienced a similar amount of unpleasant affect when witnessing the other’s pain on either hand. In other words, there was no transfer of the first-hand placebo analgesia effect, neither to empathy for pain nor to one’s own unpleasantness.

#### 3.1.2 Post hoc analyses

Complementing the above results, the Bayesian paired *t*-tests using self- and other-related pain as well as unpleasantness ratings showed very strong evidence for a placebo analgesia effect in first-hand pain (BF_10_ = 3.15 × 10^9^), but strong evidence against such an effect in pain empathy (analysis A3 in Figure 2). The latter was visible in the other-related pain ratings where the null hypothesis was found to be approximately eight times more likely than the alternative hypothesis in our sample (BF_01_ = 8.47). For unpleasantness ratings, the null hypothesis was found to be approximately seven times more likely than the alternative hypothesis (BF_01_ = 6.80). In sum, the behavioral results suggested a strong placebo analgesia effect for self-related pain, localized to participants’ right hands, but no transfer of this effect to other-related pain or self-experienced unpleasantness.

### 3.2 fMRI results

#### 3.2.1 Manipulation checks

We conducted preregistered three manipulation checks to evaluate the (i) validity of our design, (ii) success of the first-hand placebo analgesia induction and (iii) existence of overlapping activation for self- and other-related pain (see Figure 5).

**Figure 5.**
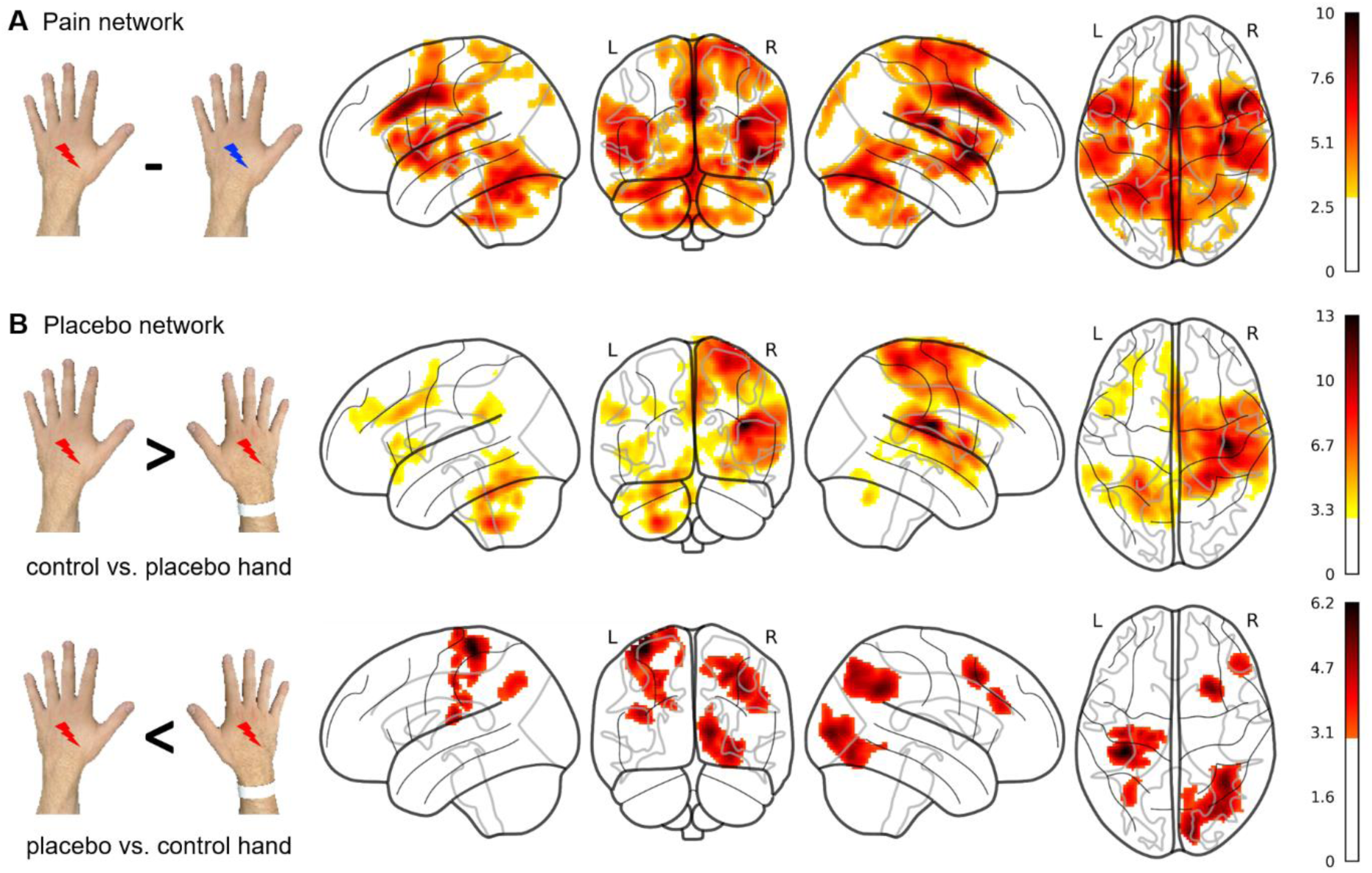
Preregistered manipulation checks of the fMRI data (see also Figure 2). A) Check b1 aimed at showing the first-hand pain processing network induced by electrical stimulation and is displayed as the contrast [*self - pain* (red icon) *> self - no pain* (blue icon)] for the control hand only. Here, we observed increased activity in both affective-motivational and sensory-discriminative pain processing regions. B) Check b2 aimed at evaluating the existence of a placebo analgesia network and is displayed using the contrasts *[self - pain - control hand > self - pain - placebo hand]* and *[self - pain - placebo hand > self - pain - control hand]* (results of this check are reported in the text using small volume correction (SVC) on specific regions). There, we observed the typical placebo network and initial evidence for a first-hand localized placebo effect. Check b3 was done using SVC and is therefore not displayed in here but used the conjunction *[self - pain > self - no pain] ∩ [other - pain > other - no pain]*. This revealed increased activation in left AI when using only the control hand, and bilateral AI as well as aMCC when averaging over both hands. All statistical activation maps in the figure are displayed whole brain, FWE-corrected at *p* < .05 cluster correction (*k* = 188) and an initial cluster-forming threshold of *p* < .001 uncorrected. L = left hemisphere; R = right hemisphere.

For check (i), the contrast *[self - pain > self - no pain]* for the control hand revealed increased hemodynamic activity in three major clusters encompassing, among others, ACC, MCC, bilateral insula, bilateral S2, thalamus and cerebellum (manipulation check b1 in Figure 2; see A.7 and Table A3 in Supplement A) whole brain, *p* < .05 FWE-corrected at cluster level). These results showed that typical sensory-discriminative and affective-motivational areas of first-hand pain processing were activated by our task.

For check (ii), we evaluated the contrasts [*self - pain - control hand > self - pain - placebo hand]* and [*self - pain - placebo hand > self - pain - control hand]* (manipulation check b2 in Figure 2; see A.7 and Table A4 in Supplement A; SVC, *p* < .05 FWE-corrected at peak-level). We found increased hemodynamic activity in right S2, right posterior insula, bilateral dACC, bilateral AI and thalamus when participants received painful stimulation on the left control compared to the right placebo hand. In the opposite contrast, we observed increased activity in right DLPFC and left S2 for the right placebo compared to the left control hand. As whole brain results of these two contrasts encompassed multiple additional regions, these results are reported in Table A5 in Supplement A.

For check (iii), the conjunction *[(self pain > self no pain) ∩ (other pain > other no pain)]* using contrasts of the control hand revealed increased activation in left AI (manipulation check b3 in Figure 2; SVC, *p* < .05 FWE-corrected at peak-level). When averaging over both hands, we observed increased hemodynamic activity in bilateral AI and aMCC.

#### 3.2.2 Preregistered analyses

After having verified the overall validity and effectiveness of the experimental paradigm as well as the placebo induction procedures, we went on to test our main hypothesis for a transfer of the first-hand placebo analgesia effect to empathy for pain using a ROI approach (see Table 1 for an overview of all paired *t*-tests). To this end, we extracted parameter estimates of three affective (bilateral AI, aMCC) and four somatosensory ROIs (bilateral S1 and S2). We first calculated an ANOVA pooling the activation of all seven ROIs and then calculated separate ANOVAs and planned comparisons for each ROI to evaluate the first-hand placebo effect, its transfer to pain empathy, and to compare the effects for self- and other-related stimulation.

**Table 1.** Main ROI analyses testing for self- and other-related placebo effects in affective and somatosensory brain regions, as well as for differences between self- and other-related effects via paired t-tests.

|  |  | self |  |  | other |  |  | self vs. other |  |  |
| --- | --- | --- | --- | --- | --- | --- | --- | --- | --- | --- |
|  |  | *t*(44) | *p* | *d<sub>z</sub>* | *t*(44) | *p* | *d<sub>z</sub>* | *t*(44) | *p* | *d<sub>z</sub>* |
| **B1** | pooled ROIs | 0.66 | .256 | 0.09 | 0.23 | .409 | 0.03 | -0.27 | .394 | -0.04 |
| **B2** | left AI | 1.60 | **.059<sup>†</sup>** | 0.24 | 0.67 | .254 | 0.10 | -0.63 | .267 | 0.09 |
|  | right AI | 0.41 | .341 | 0.06 | -0.54 | .298 | 0.08 | -0.57 | .285 | 0.09 |
|  | aMCC | -0.07 | .473 | 0.01 | 0.41 | .341 | 0.06 | 0.32 | .376 | 0.05 |
|  | left S1 | -0.20 | .423 | 0.03 | 0.42 | .338 | 0.06 | 0.39 | .349 | 0.06 |
|  | right S1 | 0.28 | .391 | 0.04 | 0.35 | .366 | 0.05 | 0.04 | .483 | 0.006 |
|  | left S2 | -1.87 | **.034<sup>\*</sup>** | 0.28 | -0.95 | .174 | 0.14 | 0.57 | .285 | 0.09 |
|  | right S2 | 3.27 | **.001<sup>\*</sup>** | 0.49 | 0.37 | .355 | 0.06 | -1.87 | **.034<sup>\*</sup>** | 0.28 |
| **B3** | IS1 vs. rS1 | 0.88 | .385 | 0.13 | -0.74 | .465 | 0.11 | 1.07 | .292 | 0.16 |
|  | IS2 vs. rS2 | -4.30 | **<.001<sup>\*</sup>** | 0.64 | -0.75 | .455 | 0.11 | -2.85 | **.006<sup>\*</sup>** | 0.42 |

Note. Planned comparisons for the region of interest (ROI) analyses (analyses B1 and B2 in Figure 2) to evaluate the first-hand placebo effect (self), its transfer to pain empathy (other) and to compare the effects for self- and other-related stimulation (self vs. other). Furthermore, analysis B3 (here and in Figure 2) directly compared activity in right vs. left S1, and right vs. left S2, related only to stimulation of the contralateral hand. *p* values for preregistered analyses B1 and B2 are reported one-tailed, and two-tailed for analysis B3. AI = anterior insula; aMCC = anterior midcingulate cortex; r/l S1 = right/left primary somatosensory cortex; r/l S2 = right/left secondary somatosensory cortex; *t*(degrees of freedom); *d<sub>z</sub>* = Cohen's *d*; <sup>†</sup> = trend; \* *p* < .05.

The pooled ANOVA (analysis B1 in Figure 2) showed significant main effects of target, hand, intensity and ROI, a significant target x intensity interaction, as well as all interactions involving the factor ROI (except for the four way interaction target x hand x intensity x ROI, see Table A6 in Supplement A). When comparing brain activity related to the placebo vs. the control hand encompassing pooled activation of all ROIs, we found no significant differences for self-related or other-related stimulation (self: *M*_*diff*_ = 0.212, 95% CI_meandiff_ [-0.44, 0.86]; other: *M*_*diff*_ = 0.072, 95% CI_meandiff_ [-0.55, 0.70]). The magnitudes of these effects were indistinguishable between self and other (self vs. other: *M*_*diff*_ = -0.139, 95% CI_meandiff_ [-1.18, 0.90]). The absence of effects in the pooled ANOVA might be explained by differential, inhomogeneous effects in the seven ROIs. As preregistered, and due to a significant main effect of ROI as well as significant interactions with the factor ROI, we went on to calculate single ANOVAs and complementary *t*-tests for each ROI separately (analysis B2 in Figure 2).

The separate ROI analyses of the three affective regions revealed a trend in left AI for self-but not for other-related stimulation, with the control hand showing slightly increased activation compared to the placebo hand (self: *M*_*diff*_ = 0.825, 95% CI_meandiff_ [-0.22, 1.87]; other: *M*_*diff*_ = 0.315, 95% CI_meandiff_ [-0.63, 1.26]). We found no significant differences between placebo and control hand in right AI or aMCC, neither for self-nor other-related stimulation (right AI, self: *M*_*diff*_ = 0.200, 95% CI_meandiff_ [-0.78, 1.18], other: *M*_*diff*_ = 0.211, 95% CI_meandiff_ [-1.00, 0.58]; aMCC, self: *M*_*diff*_ = 0.034, 95% CI_meandiff_ [-1.04, 0.97], other: *M*_*diff*_ = 0.218, 95% CI_meandiff_ [-0.85, 1.29]). The magnitudes of these effects were indistinguishable between self and other (self vs. other, left AI: *M*_*diff*_ = -0.510, 95% CI_meandiff_ [-2.15, 1.13]; right AI: *M*_*diff*_ = -0.411, 95% CI_meandiff_ [-1.86, 1.04]; aMCC: *M*_*diff*_ = 0.253, 95% CI_meandiff_ [-1.35, 1.86]; see Figure 6 here and Table A7-A9 in the Supplement).

**Figure 6.**
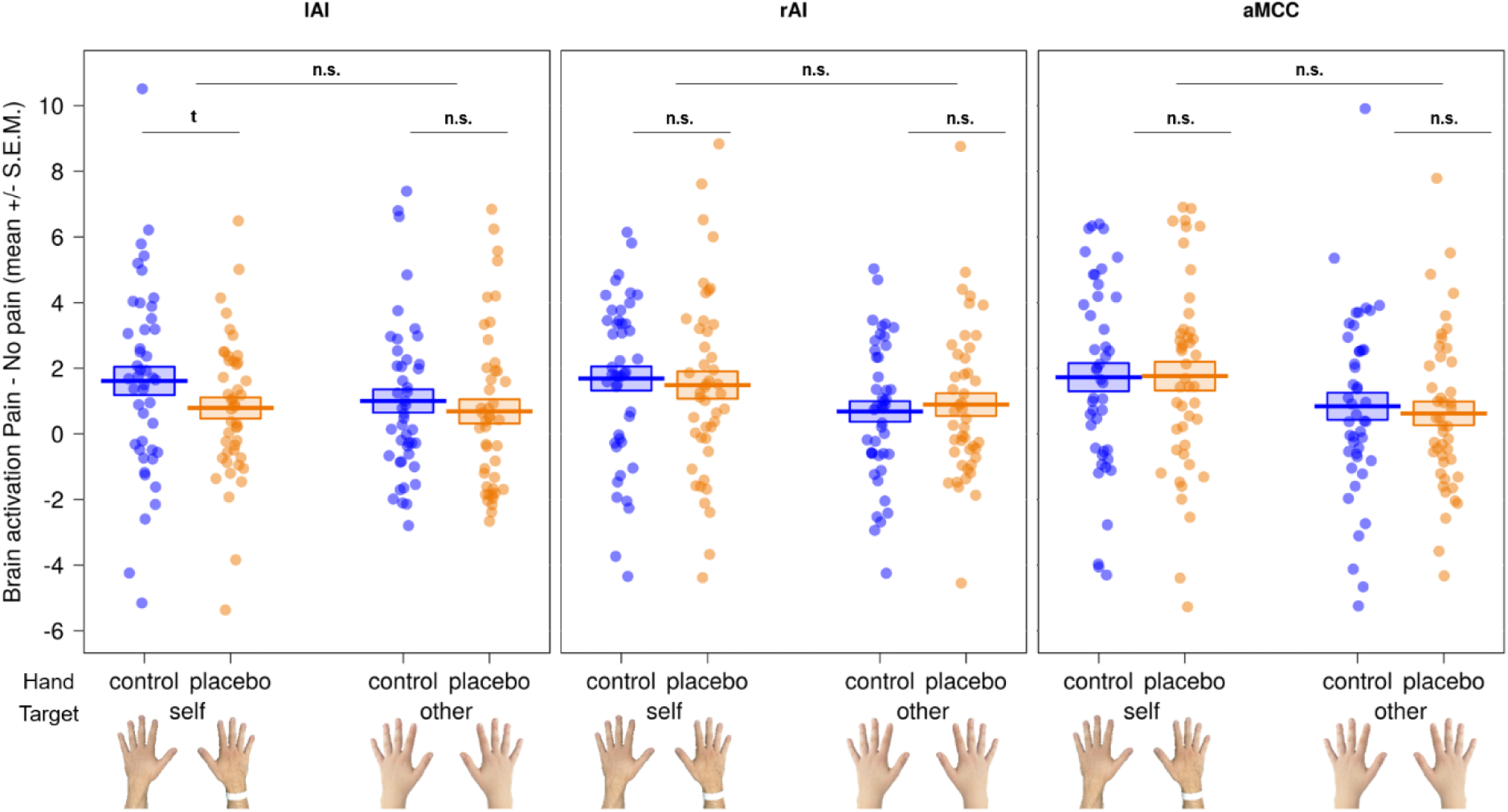
Paired comparisons of the region of interest (ROI) results for affective brain regions. Results in left anterior insula (lAI), right anterior insula (rAI) and anterior midcingulate cortex (aMCC) revealed no modulation in the three affective ROIs for self or other by the placebo manipulation. In other words, both hands led to similar hemodynamic activity in each ROI. We did find a trend (t) of *p* = .059 one-tailed in left AI, with increased activity during stimulation of the control hand in the self condition. n.s. = not significant; S.E.M. = standard error of the mean.

The four somatosensory ROIs showed differential results for S1 and S2. The planned comparisons in S1 revealed no differences between the hands, neither for self-nor other-related stimulation (left S1, self: *M*_*diff*_ = -0.091, 95% CI_meandiff_ -1.03, 0.85], other: *M*_*diff*_ = 0.181, 95% CI_meandiff_ [-0.68, 1.04]; right S1: self: *M*_*diff*_ = 0.116, 95% CI_meandiff_ [-0.73, 0.96], other: *M*_*diff*_ = -0.142, 95% CI_meandiff_ [-0.69, 0.97]). The magnitudes of these effects were indistinguishable between self and other (self vs. other, left S1: *M*_*diff*_ = 0.271, 95% CI_meandiff_ [-1.13, 1.67]; right S1: *M*_*diff*_ = 0.026, 95% CI_meandiff_ [-1.22, 1.27]).

For left and right S2, however, we observed significant differences between placebo and control hand for self-but not for other-related stimulation (left S2, self: *M*_*diff*_ = -0.461, 95% CI_meandiff_ [-0.96, 0.04], other: *M*_*diff*_ = -0.262, 95% CI_meandiff_ [-0.82, 0.30]; right S2, self: *M*_*diff*_ = 0.928, 95% CI_meandiff_ [0.36, 1.50]; other: *M*_*diff*_ = 0.124, 95% CI_meandiff_ [-0.54, 0.79]). Interestingly, activity in left S2 was significantly higher for the contralateral placebo hand while this effect was reversed in right S2 (higher activity for contralateral control hand; see panel A in Figure 7 here and Tables A10-A11 in Supplement A for full ANOVAs). The magnitudes of these effects were not different between self and other in left S2, but significantly different in right S2, where the difference between the two hands was higher for in the self condition (self vs. other, left S2: *M*_*diff*_ = 0.199, 95% CI_meandiff_ [-0.50, 0.90]; right S2: *M*_*diff*_ = -0.804, 95% CI_meandiff_ [-1.67, 0.06]; see panel B in Figure 7 here and Tables A12-A13 in Supplement A for full ANOVAs).

**Figure 7.**
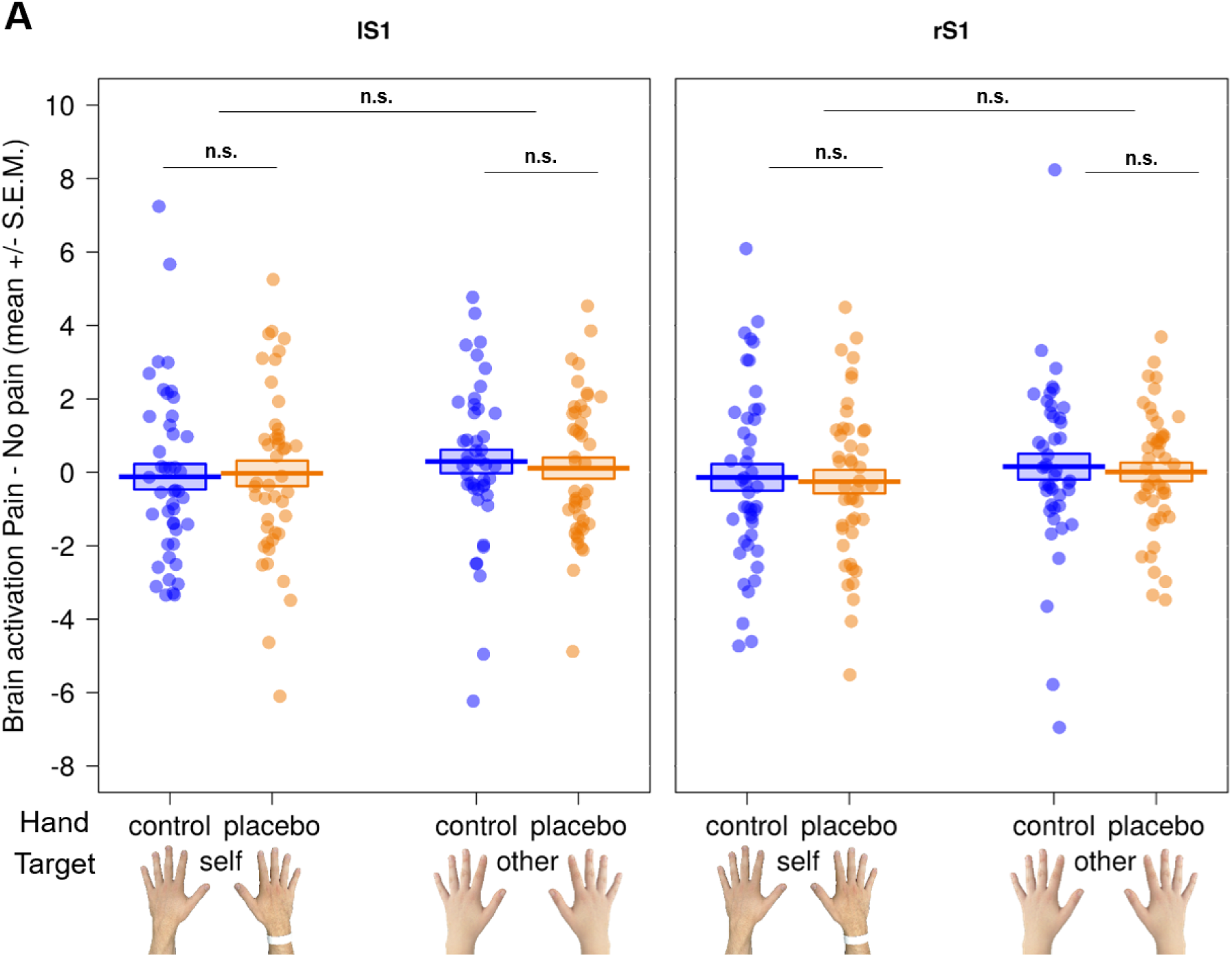
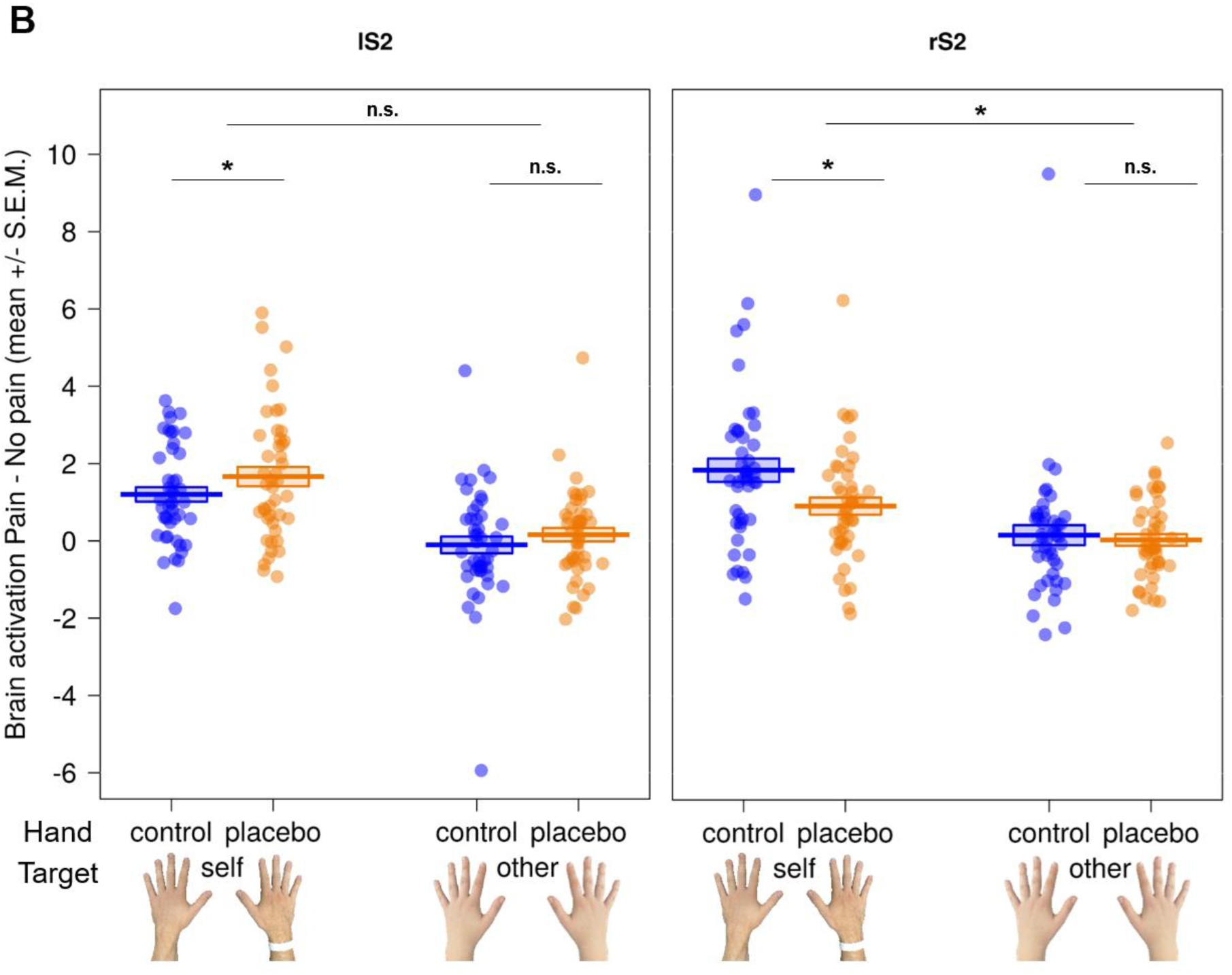
Paired comparisons of the region of interest (ROI) results for somatosensory brain regions. A) Results in left (lS1) and right (rS1) primary somatosensory cortex revealed no evidence for a modulation by the placebo manipulation in either hemisphere, neither for self nor other. B) Results in left (lS2) and right (rS2) secondary somatosensory cortex showed differential effects: In the self condition, hemodynamic activity was significantly increased in lS2 for the contralateral placebo hand, while activity was higher in rS2 for the contralateral control hand. Generally, we found no significant differences regarding other-related stimulation, but the first-hand placebo effect in lS2 was significantly stronger than its other-related counterpart. \**p* < .05; n.s. = not significant; S.E.M. = standard error of the mean.

#### 3.2.3 Post hoc analyses

Lastly, to gather more evidence for a localized placebo effect, we compared activation in each hemisphere related only to the contralateral hand with each other, for self- and other-related stimulation, respectively (analysis B3 in Figure 2; see Table 1 and Figure 8).

**Figure 8.**
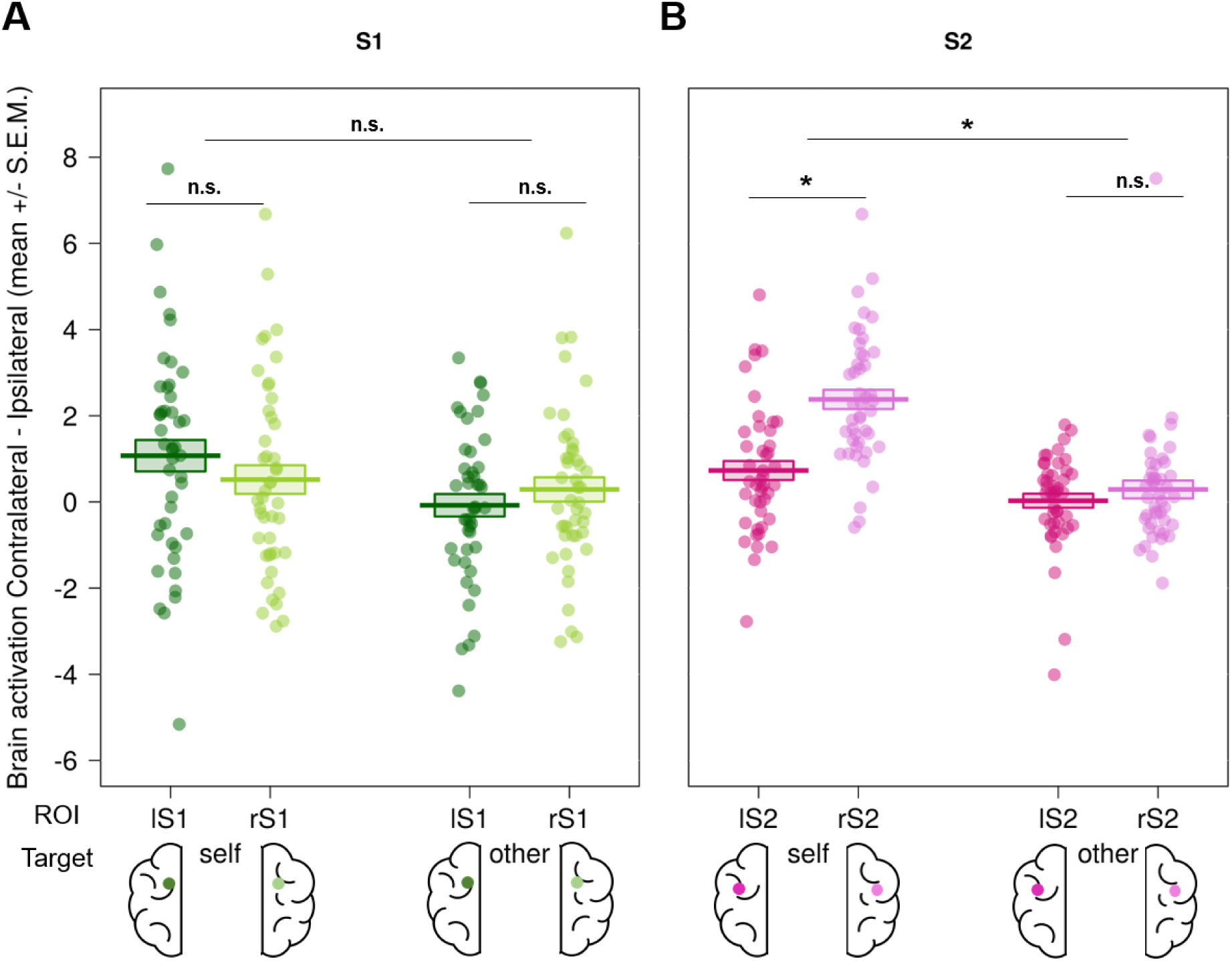
Evidence for a first-hand localized placebo analgesia effect in secondary somatosensory cortex (S2). We compared activity for each hemisphere related only to the contralateral hand with each other, i.e. activation in right primary somatosensory cortex (S1) during stimulation of left (contralateral) control hand vs. activation in left S1 during stimulation of the right (contralateral) placebo hand, and the same for secondary somatosensory cortex (S2). This was done separately for self- and other-related stimulation. A) For S1, we found no evidence for a modulation by the placebo manipulation. B) For S2, we observed increased activity in right compared to left S2 in the self condition. In general, we did not find a difference between hemispheres for the other-condition. The difference in S2 for first-hand pain was significantly stronger than its other-related counterpart. \**p* < .05; n.s. = not significant; S.E.M. = standard error of the mean.

Mirroring the ROI results above, we found differential results for S1 and S2. Regarding S1, there was no difference in brain activation between control and placebo hand for self- or other-related stimulation (self: *M*_*diff*_ = 0.553, 95% CI_meandiff_ [-0.72, 1.82]; other: *M*_*diff*_ = -0.37, 95% CI_meandiff_ [-1.37, 0.64]). The magnitude of these effects did not differ between self and other (self vs. other: *M*_*diff*_ = 0.92, 95% CI_meandiff_ [-0.82, 2.66]).

Regarding S2, we found a significant difference in brain activation between control and placebo hand for self-related pain, with the right S2 contralateral to the control hand showing increased activation compared to the left S2 contralateral to placebo hand (rS2 vs. lS2 for self: *M*_*diff*_ = -1.65, 95% CI_meandiff_ [-2.42, -0.88]); *M*_*rS2*_ ± *SD* = 2.38 ± 1.49, *M*_*lS2*_ ± *SD* = 0.73 ± 1.48). In other words, stimulation of the control hand produced significantly greater contralateral S2 activation than stimulation of the placebo hand. Regarding other-related stimulation, we did not find a difference between the right and left S2 (rS2 vs. lS2 for other: *M*_*diff*_ = -0.26, 95% CI_meandiff_ [-0.96, 0.44]; *M*_*rS2*_ ± *SD* = 0.29 ± 1.39, *M*_*lS2*_ ± *SD* = 0.03 ± 1.09). Comparing these two effects between self and other showed a significant difference, i.e. evidence for a placebo effect for the self, but not for the other (self vs. other: *M*_*diff*_ = -1.39, 95% CI_meandiff_ [-2.37, -0.41], *M*_*self*_ ± *SD* = 1.65 ± 2.57, *M*_*other*_ ± *SD* = 0.26 ± 2.32).

In sum, we replicated previous results regarding shared activations, as we observed an overlap of affective brain regions for self- and other-related stimulation. In line with the behavioral results, the fMRI results suggested that we successfully induced a localized first-hand placebo analgesia effect in the right hand of our participants, visible in increased brain activity related to the left control hand in contralateral S2. This effect, however, did not transfer to empathy for another’s pain, as we did not observe modulation of brain activity in S2 (or S1) by the placebo in the empathy condition.

## 4 Discussion

In this preregistered study, we addressed the debated question what role somatosensory aspects of the first-hand pain experience play during empathizing with someone else in pain. In particular, we wanted to pinpoint whether, when we witness another’s pain, sharing their somatosensory representations plays a similar causal role as previously shown for affective representations (e.g. Rütgen, Seidel, Silani, et al., 2015). To test this question, we induced localized placebo analgesia on the right hand of 45 participants by means of a placebo gel (with the left hand acting as a control). We then measured brain activity with fMRI during a pain task tailored towards observing possible involvement of the sensory-discriminative component of empathic pain processing. While our findings indicated both behavioral and fMRI evidence for a robust first-hand, localized placebo analgesia effect, we did not observe a transfer of this effect to empathy for pain. We thus found no causal evidence for the involvement of the sensory-discriminative component in the processing of empathic pain.

Regarding pain empathy, our findings replicated the well-documented overlap between first-hand and empathy for pain in bilateral AI and aMCC, as reported extensively in previous studies (Corradi-Dell’Acqua et al., 2011; Lamm et al., 2011 for a meta-analysis; Ochsner et al., 2008; Singer et al., 2004; Zaki et al., 2016 for a review). Moreover, the other-related pain and self-experienced unpleasantness ratings indicated that participants engaged in the task and felt empathy for the other person. Thus, participants not only correctly evaluated the pain of the other person, but also showed an empathic response. However, despite these findings, we did not observe a localized transfer of the first-hand placebo analgesia effect to empathy. Behaviorally, participants showed no reduction in empathy ratings for the placebo hand. This lack of inference-statistical significance was further corroborated by much smaller effect sizes, and strong evidence against a transfer of this effect to empathy in the Bayesian analyses. Analysis of the fMRI data directly mirrored these results, as we did not observe any differences in other-related brain activation between the two hands.

Although we did not observe a transfer of the placebo analgesia effect to empathy for pain, our behavioral results regarding self pain showed a strong, localized placebo analgesia effect, as evidenced by significantly reduced pain ratings for the placebo hand compared to the control hand, a large effect size and very strong evidence for this effect in the Bayesian analyses. This was expected, as our sample criteria excluded nonresponders to the placebo manipulation. We corroborated this finding on the neural level by showing increased activation in right S2 during stimulation of the contralateral control hand compared to the placebo hand, while this effect was reversed in left S2, which indicated stronger activation for the contralateral placebo hand. These results mirror studies finding a contralateral bias in somatosensory brain areas (Bingel et al., 2003; Coghill et al., 1999; Ploner et al., 1999; Singer et al., 2004; Symonds et al., 2006). When specifically comparing contralateral activation related to each hand with each other, we found further evidence in S2, with stronger activation in right S2 (related to the contralateral control hand) compared to left S2 (related to the contralateral placebo hand). Furthermore, we observed increased hemodynamic activity in affective brain areas and contralateral S2 in the control hand compared to the placebo hand, as well as increased activity in DLPFC and contralateral S2 during stimulation of the placebo hand compared to the control hand. These results replicate the typical placebo analgesia network reported in prior studies using similar local (Eippert et al., 2009; Geuter et al., 2013; Schafer et al., 2015; Schenk et al., 2014; Tinnermann et al., 2017), or global placebo analgesia inductions (Mischkowski et al., 2016; Rütgen, Seidel, Riečanský, et al., 2015; Rütgen, Seidel, Silani, et al., 2015; see Colloca et al., 2013; Meissner et al., 2011; Wager et al., 2011 for reviews). Together, these results demonstrate the successful induction of a first-hand, localized placebo analgesia effect on the behavioral and neural level (for S2). Interestingly, we found no evidence for such an effect in our chosen ROIs of right and left S1 representing the “hand areas”. S1 has often been implicated in the processing of stimulation in general (Ploner et al., 2000) and our design subtracted non-painful stimulation to control for unspecific touch-related activation. In fact, we did not observe any activation in the whole brain contrast [*self - pain > self - no pain]* for both hands in the area we selected for our ROI analysis (Bingel et al., 2004), but instead observed activation in a different, more dorsomedial cluster.

Although we found increased brain activity in affective-motivational brain areas for the control compared to the placebo hand on a whole brain level, our ROI analyses did not show any modulation by the placebo manipulation during self-related stimulation (except for a trend in right AI showing increased activity for the control hand, consistent with our predictions). This may seem contradictory to what was reported by Rütgen, Seidel, Silani, et al. (2015). However, it should be noted that in that study, two groups with either placebo analgesia or control were compared, while the current design made comparisons within participants who all underwent placebo analgesia, with a specific focus on somatosensory aspects of the pain experience. Moreover, we employed a localized compared to a generalized placebo analgesia induction, which may also have influenced the affective-motivational component of first-hand pain. Thus, we cannot draw any conclusions about the here absent modulation of affective regions during self-experienced pain.

To answer our research question, we documented clear evidence for a localized placebo effect in first-hand pain but find no evidence for a transfer of this effect to empathy for pain, and thus no evidence for somatosensory sharing. As these results were contrary to our preregistered predictions, we now discuss why this could have been the case, highlighting strengths and possible limitations. First of all, we preregistered our design and procedure as well as most of the planned analyses prior to data collection and clearly distinguish those from post hoc analyses, thereby minimizing the possibility of false-positives and *p*-hacking (Crüwell et al., 2019; Nosek et al., 2018; Wicherts et al., 2016). Furthermore, we purposefully used a within-subjects design and an *a priori* power analysis to maximize sensitivity and the possibility of finding an effect (Beck, 2013; Charness et al., 2012). In contrast to previous studies, our design was specifically tailored to being able to observe possible somatosensory modulation. Our findings are further supported by observation of the typical pain processing network during first-hand electrical stimulation, demonstrating validity of our pain paradigm (Morton et al., 2016; Xu et al., 2020). These points strongly speak for the validity of our design and additionally bolster the interpretability of our results.

Although we specifically targeted somatosensory pain processing, participants might still have focused more on the generalized, affective consequences of the other’s pain instead of processing its localized, somatosensory consequences. This would be in line with results of an ERP study by Rütgen, Seidel, Riečanský, et al. (2015), who did not find effects of placebo analgesia on both anticipation and delivery phases in the ERP components P1 and N1 or on non-painful control stimulation (for both self- and other-related conditions). While P1 is an occipital ERP component that has been shown to index an early stage of low-level visual processing and was also linked to top-down attentional processes, the visual N1 component has been associated with attention and discrimination processes (Couperus & Mangun, 2010; Slagter et al., 2016). Due to these results, the authors argued that it is unlikely that placebo analgesia changed general aspects of sensory perception or attention in their study, but targeted affective aspects of the empathic pain experience. This might suggest that somatosensory-related processes are only or more strongly recruited by the first-hand pain experience and, therefore, do not play a strong role in empathic sharing (Decety, 2010; Jackson et al., 2006; Krishnan et al., 2016; Rütgen, Seidel, Riečanský, et al., 2015; Rütgen, Seidel, Silani, et al., 2015). Sharing the pain of others could therefore also be possible in the absence of first-hand nociception, which is important in the context of shared representations between one’s own and empathic pain experiences. Our results indicate that previously found empathy-related activations of sensorimotor processes do not necessarily indicate a specific sharing of another’s pain in one’s own pain processing system. In fact, a meta-analysis by Lamm et al. (2011) proposed that previously reported somatosensory activation during empathy for pain could reflect “rather unspecific co-activation elicited by the display of body parts being touched rather than a specific matching of the other’s somatosensory and nociceptive state” (p. 2499), as this activation was observed bilaterally and for painful and non-painful stimulation in the meta-analysis (but see also Keysers et al., 2010). In line with this argumentation, Keysers et al. (2004) found S2 (but not S1) to be active during both first-hand and empathy for touch, which matches our results on the first-hand placebo analgesia effect being represented only in S2.

Sharing of another individual’s pain might be especially focused on its affective aspects, when a fast and effective processing of the situation does not necessarily require specific somatosensory-related knowledge of pain. In our task, the perception of how unpleasant or aversive the stimulation was for the other, i.e. a general processing of that pain and its related affective consequences, might have been a more relevant dimension than the exact location of that pain (i.e. the hand). Future studies may thus want to differentiate between situations when observing another person in pain is merely related to affective sharing per se, versus a prompt for specific knowledge about another’s pain, such as when specific helping behavior is required. For instance, it may make a difference if participants are only asked to “resonate” with the pain of others without any specific request, as in our study, compared to a setup simulating e.g. the work of medical professions, where it does not suffice to resonate with the affective response but where the exact source of the pain is of higher relevance. A recent review suggested that sensorimotor activations to another’s pain could also reflect “activation of defensive responses in agreement with the goal of pain”, in order to protect the body from external harm (Riečanský & Lamm, 2019, p. 970). Those responses could thus be seen as less relevant, when the situation is known to be unpleasant and aversive but does not require helping behavior. This may also explain the discrepancy of our findings with a recent study finding a causal role of S1 in driving prosocial behavior (Gallo et al., 2018). In addition to this reasoning, previous studies showing a role of sensorimotor or somatosensory brain regions in pain empathy used a) salient video stimuli depicting painful needle injections into body parts and/or b) different setups and instructions, specifically prompting participants to reason about the sensory consequences of the stimulation and direct their attention to the specific, affected body part (Avenanti et al., 2005, 2006; Bufalari et al., 2007; Lamm et al., 2007; Motoyama et al., 2017). Despite our findings, i.e. an absence of evidence for somatosensory sharing, we therefore cannot completely rule out the possibility of still having missed somatosensory involvement with our design, since most of the studies reporting somatosensory brain activation in response to empathic processing used picture-based tasks where explicit images of limbs in painful situations are shown, while we used electrical stimulation in the present study (Lamm et al., 2011 for a meta-analysis; Xiang et al., 2018 for a review). We are currently investigating this possibility in a separate study employing a typical picture-based paradigm. However, finding complementary results in both behavior and brain responses and further evidence in our post hoc analyses, we are confident in our conclusion that the somatosensory component of pain does not play a causal role in pain empathy, in the present design.

## 5 Conclusion

Our findings suggest a robust localized placebo analgesia effect for first-hand pain, but no evidence for a role of the sensory-discriminative component in empathic sharing. Nevertheless, we observed shared brain activations between first-hand and empathy for pain in the affective-motivational component. Using a causal-experimental manipulation and a tailored design, empathy for another person was not influenced by a localized pain reduction in a specific body part, thereby not confirming our preregistered predictions. These insights are important when trying to characterize the magnitude of influence that our own pain experience has on our ability to empathize and suggest that empathy for pain, at least when investigated with the type of paradigms used here and previously, may rely more on sharing of another’s affective, compared to their somatosensory state.

## 6 Acknowledgements

We thank Ronald Sladky for input on a final draft of the preregistration and Paul Forbes for feedback on the final draft of the manuscript. We would also like to thank the two master students Fabian Franken and Anna Köstler who worked on this project as well as the numerous interns and confederates helping with data collection.

## 7 Author contributions

**Helena Hartmann:** Conceptualization, Data curation, Formal analysis, Funding acquisition, Investigation, Methodology, Software, Visualization, Writing - original draft, Writing - review & editing. **Markus Rütgen:** Formal analysis, Methodology, Supervision, Writing - original draft, Writing - review & editing. **Federica Riva:** Formal analysis, Supervision, Writing - original draft, Writing - review & editing. **Claus Lamm:** Conceptualization, Formal analysis, Funding acquisition, Methodology, Project administration, Resources, Supervision, Writing - original draft, Writing - review & editing.

