## Supplements A and B for "Another’s pain in my brain: No evidence that placebo analgesia affects the sensory-discriminative component in empathy for pain"

### Supplement A

#### A.1 Sample

Exclusion criteria were past or present enrollment in academic studies including psychology, pharmaceutics or (veterinary, dental, human, etc.) medicine (due to prior knowledge about deceptive elements of the study setup), any neurological or psychiatric conditions, past or present self-injurious behavior, past or present medical conditions regarding the hands or the skin on the hands that might interfere with current pain sensitivity (e.g. numbness, chronic pain, trauma or injuries, or long-term pain therapy), past or present substance abuse (alcohol or drugs), the intake of psychopharmacological medication (besides oral contraceptives) within the last three months, left-handedness or no handedness preference, and a weakness in distinguishing right from left. Handedness was assessed with a combination of two handedness questionnaires (Büsch et al., 2009; Oldfield, 1971). As suggested by Tran, Stieger, & Voracek (2014), a laterality quotient (LQ; going from -100 = strongly left-handed to +100 = strongly right-handed) was calculated out of the ten most selective items of both questionnaires, and only individuals with LQs of 72 or higher were invited to the study. A weakness in distinguishing right from left was operationalized as answering the question “How often do you mix up left and right in your daily life (e.g. while making a turn with the car)?” with either ‘sometimes’, ‘often’ or ‘always’. We tested each participant twice for the contraindications regarding MRI scanning, once via the online screening questionnaire and once face-to-face at the outset of the scanning session. Dropouts, including placebo analgesia nonresponders, were replaced until the calculated sample size of 45 participants was reached. We excluded 20 participants (25.6 %) for not responding to the placebo manipulation, seven due to technical malfunctioning of the pain stimulator, five for inconsistent ratings (e.g. similar ratings for painful and non-painful conditions independent from the placebo manipulation) or extensive movement and one due to a spontaneously found abnormality in the brain, adding up to a total of 78 recruited participants.

#### A.2 Nonresponder identification

Nonresponders in regard to the placebo manipulation were determined using four exclusion criteria. First, any verbally expressed doubts regarding the study setup were recorded during and after the session and followed up on. In case of vague doubts, participants were asked to specify their doubts as much as possible. Second, we assessed belief scores about the effectiveness of the medication to decrease pain at three time-points during the session (after the medical cover applied the gel = pre-conditioning, after the conditioning procedure = post-conditioning, and after the completion of all tasks = post-session). Low beliefs in general (pre- and post-conditioning ratings each on a continuous visual-analogue-scale from 0-10 cm, sum of both ratings < 6.66 cm) and strong decreases between the first and the second time-point (pre- minus post-conditioning ratings > 3.33 cm) indicated a lack of responding. Third, we took the number of needed conditioning trials into account. If participants responded with a rating higher than 5 to the conditioning stimulus on the right hand (given at a medium intensity of 4) and/or with a rating of 5 or lower to the stimulus on the left hand (given at a high intensity of 7), the trial was deemed unsuccessful and repeated. It should be noted here that none of the tested participants needed more than three conditioning trials. Finally, due to the within-subjects design of this study, we were able to directly compare the first-hand pain ratings of the left and right hand and use them as an additional exclusion criterion.

#### A.3 Procedure

To ensure equal conditions for all participants and to strengthen the cover story, we instructed participants to refrain from alcohol, drugs or medication-intake 24 hours before the session as well as from intake of food or any other drink but water one hour before the session. Participants were told that the gel was already well established regarding its effectiveness, but we were interested in its neural representation.

For the placebo induction we used a combination of verbal suggestions and classical conditioning techniques. First, the medical cover (who was either male or female) did a short medical screening including a (pseudo) drug-test to increase believability of the cover story, after which he/she explained the pain-reducing effects of the “medication” and gave further information on the duration of action, use and side-effects. The “medication” was presented to the participants as a “potent, local anesthetic” with strong pain-reducing effects only on the part of the skin it is applied to. We used the term “anesthesia” rather than “analgesia” in the instructions to increase believability, as the placebo was a topical gel. Furthermore, the “medication” was presented as being well established and legally approved since many years, as well as frequently and routinely used for chronic pain patients and in dental procedures. As a side-effect in very rare cases, dry skin and slight irritation was mentioned. To avoid decrease of beliefs due to an absence of “numbing”, participants were told that the medication blocked pain receptors pharmacologically and thus would exert no effects on normal touch sensitivity or non-painful stimulation. Participants were further told that the effectiveness of the medication would be tested after the waiting time by means of a brief pain test from the experimenters before going into the scanner. If the participant had no more questions, the medical cover first applied the placebo gel on the whole dorsum of the right hand (on which the medical cover had previously attached a white paper bracelet to visually remind the participants of their “treated” hand) and directly after that a control gel with no active ingredients on the left hand. The medical cover explained to the participants that this was routinely done to ensure equal conditions for the two hands and make them comparable for later analysis. He/she further explained that the inactive gel on the left hand represented a basis for the participant’s normal pain perception in the study (the word “placebo” was never mentioned). To compare brain activation of all participants in the group analysis later on, the placebo gel was applied on the dominant, right hand and the control gel on the left hand in all participants. Both gels contained 0.5 g Carbomer, 0.09 g TRIS, 15 g undiluted Isopropanol, 3 g Glycerol, 10 g Propylenglykol and Aqua pur. ad 100 g. The only difference was 10 g (out of 100 g) more Isopropanol in the placebo gel and 10 g basic skin cream (‘Ultrasicc’) instead of Isopropanol in the control gel to change visual and olfactory properties.

In the scanner room, the confederate was told that she should communicate with gestures during the experiment because the microphone was built inside the scanner bore. These instructions were given in front of the participant to further enhance the belief that the confederate was in fact a second participant. In general, previously used procedures were used to ensure believability of the whole setup, e.g. ensuring contact every now and then between confederate and participant, attaching fake electrodes to the confederate’s hands, conducting fake electrode tests with the confederate to check whether they were adhered to the skin correctly, and communicating with both participants equally during the scanning breaks. However, the confederate never received any electrical stimulation, covertly left the scanner room before the scanning started, stayed in the control room with the experimenters during the scanning and exited the building before the participant came out of the scanner. In case of breaks in between runs or emergencies, the confederate was secretly asked to return to her table next to the scanner as if she had been sitting there during the task. To avoid habituation of the skin where the electrodes were attached and confusion of painful and non-painful stimulation, we applied one additional electrode on each dorsum of the hand for delivery of the non-painful stimulation. All four electrodes were tested with the participant lying down by giving low intensity stimulation on each electrode to ensure optimal skin adherence. The value for non-painful stimulation calibrated outside of the scanner was therefore tested on the new electrodes and if rated too low or high, slightly adjusted for each hand to represent a subjective rating of 1 = “not painful, but perceivable”.

##### A.4 Pain task

The visual stimuli of participant’s and confederate’s hands were created with a Leica D-Lux 54 in the first session. Hands resting on a black cardboard were photographed from the top in similar lighting conditions. The confederate stimuli were created in the same way, but without the white bracelet to indicate no medication. The stimuli were then edited with Photoshop CC and Microsoft PowerPoint, removing the background and inserting target and hand cues in either red (RGB = 255/0/0, HEX = #FF0000) or blue (RGB = 0/0/255, HEX = #0000FF). The final stimuli were rescaled to 900x627 pixels for the task. During the scanning, the task was displayed on a BOLD screen 32 LCD for fMRI (https://www.crsltd.com/tools-for-functional-imaging/mr-safe-displays/boldscreen-32-lcd-for-fmri/nest/boldscreen-32-technical-specification#npm) that the participant viewed over a mirror mounted on the head coil.

##### A.5 Analyses and plotting

For our analyses and plotting in RStudio, we used the following packages (and functions): stats (shapiro.test, t.test, cor.test, aggregate), ez (ezANOVA), BayesFactor (ttestBF), plyr (ddply), dplyr (arrange), tidyr (gather, spread), reshape2 (dcast), yarrr (pirateplot) and ggplot2 (ggplot). Figures 3 and 5 were created in Python/Jupyter Notebook using the Nilearn module (functions: plot_anat, add_contours and plot_glass_brain; https://nilearn.github.io/index.html).

##### A.6 Behavioral results

In order to evaluate the strength of the subjective placebo effect in our sample, we conducted two manipulation checks. The first check was preregistered as exploratory, the second one was added post hoc to get a better impression of the subjective pain experience on each hand when asked at the end of the session. First, we investigated differences between the three ratings of the placebo gel effectiveness. Mean ratings were $M \pm SD$ = 6.64 ± 1.80 after the administration of the gel (pre-conditioning), 8.06 ± 1.27 after the conditioning procedure (post-conditioning) and 6.71 ± 2.62 at the end of the testing session (post-session; see manipulation checks a1 and a2 in Figure 2 in the main text and Figure A1 here). Significant differences emerged between the pre- and post-conditioning ratings ($t(44) = 5.91$, $p < .001$, $M_{diff} = 1.42$, 95% $CI_{meandiff}$ [0.94, 1.91]), as well as between the post-conditioning vs. post-session ratings ($t(44) = -3.80$, $p < .001$, $M_{diff} = -1.36$, 95% $CI_{meandiff}$ [-2.07, -0.64]). No significant differences were found between the pre-conditioning vs. post-session ratings ($t(44) = 0.16$, $p = 0.875$, $M_{diff} = 0.07$, 95% $CI_{meandiff}$ [-0.78, 0.92]). This indicated that effectiveness beliefs in the “medication” increased significantly from gel administration to the conditioning procedure. Furthermore, participants’ beliefs in the “medication” dropped after the completion of the task, but it did not drop lower than the initial effectiveness belief after gel administration when asked at the end of the session. Second, we evaluated the difference between average stimulation intensities felt on each hand when asked after the session and found a significant difference between the placebo ($M \pm SD = 5.08 \pm 1.76$) and the control hand ($M \pm SD = 7.01 \pm 0.84$): $t(44) = -8.11$, $p < .001$, $M_{diff} = -1.93$, 95% $CI_{meandiff}$ [-2.41, -1.45]. Participants recalled feeling less pain on their right placebo hand compared to their left control hand during the task. Both manipulation checks therefore showed evidence for a subjective localized placebo effect over the whole course of the session that was highest right before the completion of the pain task.

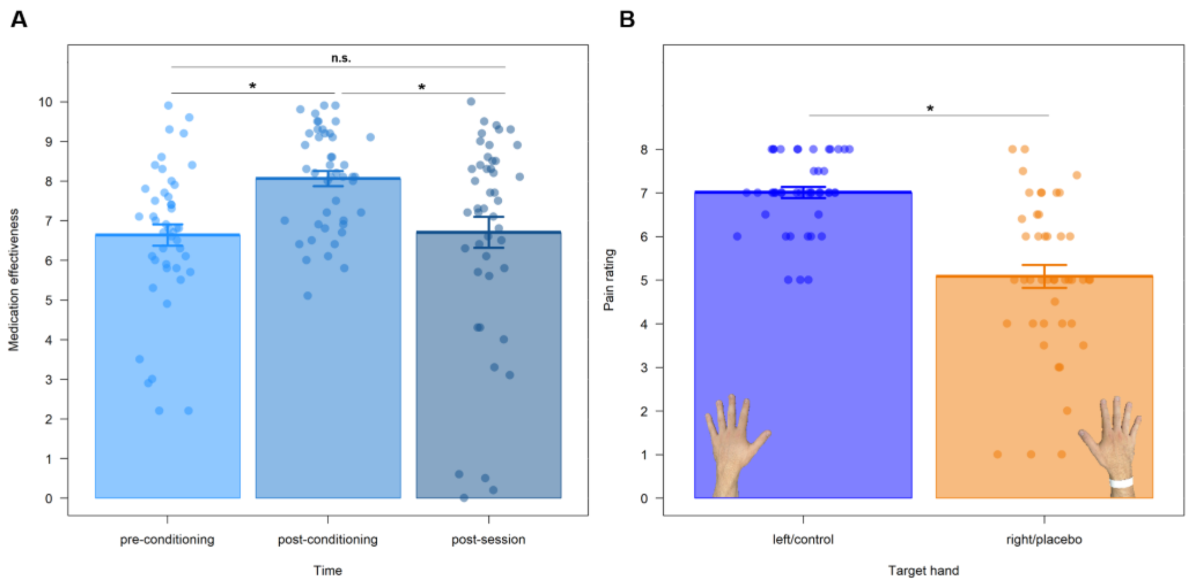


*Figure A1.* We conducted two behavioral manipulation checks evaluating the strength of the first-hand placebo effect. Displayed are means and standard errors of the mean. A) Beliefs in the effectiveness of the administered placebo gel at three time points during the experiment: after the topical application by the medical cover (pre-conditioning), after the conditioning procedure (post-conditioning) and during the debriefing at the end of the session (post-session). This revealed a significant increase of beliefs after the conditioning. Although the beliefs significantly decreased until the end of the session, they did not drop significantly lower than initial belief levels. B) Average intensity of the painful stimulation received on each hand judged at the end of the session. Here, we observed significantly lower pain ratings for the placebo compared to the control hand.

Table A1

*Behavioral results of the ANOVA using self- and other-related pain ratings*

| Effects | $F_{(1,44)}$ | $p_{(two\text{-}tailed)}$ | gen. $\eta^2$ |
|---|---|---|---|
| target | 72.15 | < .001 | 0.13 |
| hand | 18.44 | < .001 | 0.04 |
| intensity | 840.67 | < .001 | 0.84 |
| target x hand | 20.11 | < .001 | 0.04 |
| target x intensity | 9.31 | < .001 | 0.03 |
| hand x intensity | 61.54 | < .001 | 0.05 |
| target x hand x intensity | 67.58 | < .001 | 0.05 |

Table A2

*Behavioral results of the ANOVA using unpleasantness ratings*

| Effects | $F_{(1,44)}$ | $p_{(two\text{-}tailed)}$ | gen. $\eta^2$ |
|---|---|---|---|
| hand | 1.38 | .246 | 0.001 |
| intensity | 167.89 | < .001 | 0.42 |
| hand x intensity | 0.48 | .490 | < 0.001 |

#### A.7 fMRI results

Table A3

*fMRI whole brain results for manipulation check b1 (first-hand pain network).*

| Clusters and brain regions | h | *x* | *y* | *z* | *t* value | *p* value |
|---|---|---|---|---|---|---|
| **Cluster 1** (*k* = 7390) | | | | | | < .001 |
| insula | R | 38 | -14 | 20 | 10.13 | |
| insula | R | 44 | 10 | -2 | 9.62 | |
| insula | R | 38 | 4 | -10 | 8.12 | |
| **Cluster 2** (*k* = 15677) | | | | | | < .001 |
| midcingulate cortex | M | 0 | 8 | 40 | 9.54 | |
| anterior cingulate cortex | R | 2 | 24 | 28 | 9.42 | |
| midcingulate cortex | M | 0 | 16 | 34 | 9.37 | |
| **Cluster 3** (*k* = 10468) | | | | | | < .001 |
| cerebellar vermis 6 | M | 0 | -56 | -22 | 8.07 | |
| cerebellar vermis 6 | R | 2 | -76 | -18 | 7.80 | |
| cerebellum | L | -32 | -52 | -30 | 7.72 | |

*Note.* Significant clusters resulting from the contrast [*self - pain > self no pain]* in the left control hand including hemisphere h, cluster size *k*, MNI coordinates x, y, z, *t* value and *p* value (whole brain, FWE-corrected at $p < .05$, cluster level, $k > 188$). Brain regions were labelled with the Anatomy toolbox version 2.15 (Eickhoff et al., 2005).

Table A4

*fMRI SVC results for manipulation check b2 (placebo analgesia network).*

| Contrasts and brain regions | h | *k* | *x* | *y* | *z* | *t* value | *p* value |
|---|---|---|---|---|---|---|---|
| **Control hand > placebo hand** | | | | | | | |
| S2 | R | 509 | 38 | -16 | 20 | 13.37 | < .001 |
| posterior insula | R | 399 | 38 | -14 | 12 | 8.72 | < .001 |
| dACC | R | 386 | 4 | 2 | 42 | 7.22 | < .001 |
| dACC | L | 256 | -10 | 2 | 40 | 3.54 | .019 |
| anterior insula | L | 152 | -36 | 24 | 8 | 4.46 | .001 |
| anterior insula | R | 129 | 38 | 10 | 6 | 6.12 | < .001 |
| thalamus | R | 26 | 14 | -14 | 8 | 3.77 | .003 |
| **Placebo hand > control hand** | | | | | | | |
| DLPFC | R | 244 | 28 | 14 | 48 | 5.29 | < .001 |

| | | | | | | | |
|---|---|---|---|---|---|---|---|
| S2 | L | 175 | -38 | -18 | 20 | 4.90 | < .001 |

*Note.* Significant clusters resulting from the contrasts [*self - pain - control hand > self - pain - placebo hand]* and [*self - pain - placebo hand > self - pain - control hand]* including hemisphere h, cluster size *k*, MNI coordinates x, y, z, *t* value and *p* value (small volume correction (SVC), FWE-corrected at *p* < .05, peak-level). Only the highest peak is included in case of several peaks in a cluster. Brain regions were labelled with the Anatomy toolbox Version 2.15 (Eickhoff et al., 2005). S2 = secondary somatosensory cortex; aMCC = anterior midcingulate cortex; DLPFC = dorsolateral prefrontal cortex.



Table A5

*fMRI whole brain results for manipulation check b2 (placebo analgesia network).*

| Contrasts, clusters and brain regions | h | *x* | *y* | *z* | *t* value | *p* value |
|---|---|---|---|---|---|---|
| **Control hand > placebo hand** | | | | | | |
| **Cluster 1** (*k* = 16941) | | | | | | < .001 |
| rolandic operculum (OP3) | R | 38 | -16 | 20 | 13.37 | |
| postcentral gyrus (3b) | R | 26 | -40 | 68 | 10.16 | |
| precentral gyrus | R | 30 | -20 | 74 | 8.92 | |
| **Cluster 2** (*k* = 3485) | | | | | | < .001 |
| cerebellum (VIII) | L | -26 | -44 | -50 | 8.56 | |
| cerebellum (IV-V) | L | -26 | -42 | -30 | 7.21 | |
| cerebellum (VI) | L | -26 | -56 | -24 | 6.52 | |
| **Cluster 3** (*k* = 702) | | | | | | < .001 |
| rolandic operculum | L | -50 | 8 | 2 | 4.58 | |
| insula | L | -36 | 24 | 8 | 4.54 | |
| insula | L | -38 | 8 | 0 | 3.92 | |
| **Cluster 4** (*k* = 442) | | | | | | .001 |
| supramarginal gyrus | L | -52 | -38 | 28 | 5.50 | |
| supramarginal gyrus | L | -64 | -48 | 28 | 3.70 | |
| **Cluster 5** (*k* = 270) | | | | | | .011 |
| middle frontal gyrus | L | -30 | 40 | 32 | 4.02 | |
| **Placebo hand > control hand** | | | | | | |
| **Cluster 1** (*k* = 2046) | | | | | | < .001 |
| angular gyrus | R | 38 | -54 | 38 | 5.47 | |
| superior parietal lobule | R | 34 | -72 | 50 | 4.91 | |
| superior occipital gyrus | R | 36 | -74 | 48 | 4.72 | |
| **Cluster 2** (*k* = 1533) | | | | | | < .001 |
| calcarine gyrus | R | 12 | -90 | 6 | 5.40 | |
| fusiform gyrus | R | 28 | -76 | -10 | 5.08 | |
| lingual gyrus | R | 26 | -72 | -2 | 4.71 | |
| **Cluster 3** (*k* = 1466) | | | | | | < .001 |
| postcentral gyrus (4a) | L | -36 | -32 | 66 | 6.20 | |

| | | | | | | |
|---|---|---|---|---|---|---|
| paracentral lobule | L | -16 | -20 | 30 | 4.68 | |
| paracentral lobule | L | -10 | -20 | 60 | 4.67 | |
| **Cluster 4** (*k* = 439) | | | | | | .001 |
| middle frontal gyrus | R | 28 | 14 | 48 | 5.29 | |
| **Cluster 5** (*k* = 299) | | | | | | .007 |
| inferior frontal gyrus (p. triang.) | R | 46 | 34 | 24 | 4.38 | |
| middle frontal gyrus | R | 52 | 32 | 32 | 4.31 | |
| **Cluster 6** (*k* = 277) | | | | | | .010 |
| angular gyrus | L | -34 | -62 | 42 | 4.02 | |
| **Cluster 7** (*k* = 200) | | | | | | .039 |
| rolandic operculum | L | -38 | -18 | 20 | 4.90 | |

*Note.* Significant clusters resulting from the contrasts [*self - pain - control hand > self - pain - placebo hand]* and [*self - pain - placebo hand > self - pain - control hand*] including hemisphere h, cluster size *k*, MNI coordinates x, y, z, *t* value and *p* value (whole brain, FWE-corrected at *p* < .05, cluster-level, *k* > 188). Brain regions were labelled with the Anatomy toolbox Version 2.15 (Eickhoff et al., 2005).

151

Table A6

*ANOVA using pooled activation of all seven ROIs (lAI, rAI, aMCC, lS1, rS1, lS2, rS2).*

| Effects | *df* | *F* | $p_{(two\text{-}tailed)}$ | *gen.* $\eta^2$ |
|---|---|---|---|---|
| target | 1,44 | 197.03 | < .001 | 0.24 |
| hand | 1,44 | 6.48 | .014 | 0.004 |
| intensity | 1,44 | 29.58 | < .001 | 0.03 |
| roi$^{S}$ | 6,246 | 27.61 | < .001 | 0.11 |
| target x hand | 1,44 | 1.86 | .179 | 0.001 |
| target x intensity | 1,44 | 6.45 | .015 | 0.005 |
| hand x intensity | 1,44 | 0.61 | .439 | < 0.001 |
| target x roi$^{S}$ | 6,246 | 31.41 | < .001 | 0.05 |
| hand x roi$^{S}$ | 6,246 | 20.08 | < .001 | 0.01 |
| intensity x roi$^{S}$ | 6,246 | 13.86 | < .001 | 0.01 |
| target x hand x intensity | 1,44 | 0.07 | .789 | < 0.001 |
| target x hand x roi$^{S}$ | 6,246 | 12.79 | < .001 | 0.01 |
| target x intensity x roi$^{S}$ | 6,246 | 6.97 | < .001 | 0.006 |
| hand x intensity x roi$^{S}$ | 6,246 | 2.33 | .033 | 0.001 |
| target x hand x intensity x roi$^{S}$ | 6,246 | 1.01 | .419 | < 0.001 |

*Note.* Effects marked with an “S” had a significant Mauchly's test for sphericity and corresponding *p*-values are reported using Greenhouse Geisser sphericity correction. The following regions of interest were included: l/rAI = left/right anterior insula, aMCC = anterior midcingulate cortex, l/rS1 = left/right primary somatosensory cortex, l/rS2 = left/right secondary somatosensory cortex.

152

Table A7

*ANOVA of left anterior insula.*

| Effects | $F_{(1,44)}$ | $p_{(two\text{-}tailed)}$ | gen. $\eta^2$ |
|---|---|---|---|
| target | 107.71 | < .001 | 0.21 |
| hand | 7.70 | < .001 | 0.009 |
| intensity | 25.52 | .008 | 0.04 |
| target x hand | 3.12 | .084 | 0.004 |
| target x intensity | 0.91 | .345 | 0.001 |
| hand x intensity | 4.09 | .049 | 0.003 |
| target x hand x intensity | 0.39 | .533 | < 0.001 |

Table A8

*ANOVA of right anterior insula.*

| Effects | $F_{(1,44)}$ | $p_{(two\text{-}tailed)}$ | gen. $\eta^2$ |
|---|---|---|---|
| target | 195.91 | < .001 | 0.39 |
| hand | 5.51 | .023 | 0.008 |
| intensity | 32.72 | < .001 | 0.06 |
| target x hand | 2.51 | .121 | 0.003 |
| target x intensity | 4.09 | .049 | 0.006 |
| hand x intensity | 0.004 | .984 | < .001 |
| target x hand x intensity | 0.33 | .570 | < .001 |

Table A9

*ANOVA of anterior midcingulate cortex.*

| Effects | $F_{(1,44)}$ | $p_{(two\text{-}tailed)}$ | gen. $\eta^2$ |
|---|---|---|---|
| target | 138.25 | < .001 | 0.30 |
| hand | 2.84 | .099 | 0.003 |
| intensity | 27.63 | < .001 | 0.05 |
| target x hand | 2.84 | .099 | 0.003 |
| target x intensity | 5.40 | .025 | 0.008 |
| hand x intensity | 0.08 | .779 | < 0.001 |
| target x hand x intensity | 0.10 | .753 | < 0.001 |

Table A10

*ANOVA of left secondary somatosensory cortex.*

| Effects | $F_{(1,44)}$ | $p_{(two\text{-}tailed)}$ | gen. $\eta^2$ |
|---|---|---|---|
| target | 119.63 | < .001 | 0.39 |
| hand | 3.84 | .056 | 0.006 |
| intensity | 31.73 | < .001 | 0.08 |

| | | | |
|---|---|---|---|
| target x hand | 7.50 | .009 | 0.01 |
| target x intensity | 50.34 | < .001 | 0.07 |
| hand x intensity | 3.37 | .073 | 0.005 |
| target x hand x intensity | 0.33 | .569 | < 0.001 |

Table A11

*ANOVA of right secondary somatosensory cortex.*

| Effects | $F_{(1,44)}$ | $p_{(two\text{-}tailed)}$ | gen. $\eta^2$ |
|---|---|---|---|
| target | 110.67 | < .001 | 0.34 |
| hand | 135.35 | < .001 | 0.13 |
| intensity | 22.63 | < .001 | 0.06 |
| target x hand | 64.66 | < .001 | 0.08 |
| target x intensity | 42.98 | < .001 | 0.05 |
| hand x intensity | 5.66 | .022 | 0.008 |
| target x hand x intensity | 3.48 | .069 | 0.005 |

156

Table A12

*ANOVA of left primary somatosensory cortex.*

| Effects | $F_{(1,44)}$ | $p_{(two\text{-}tailed)}$ | gen. $\eta^2$ |
|---|---|---|---|
| target | 16.51 | < .001 | 0.05 |
| hand | 12.86 | < .001 | 0.02 |
| intensity | 0.18 | .678 | < 0.001 |
| target x hand | 7.54 | .008 | 0.01 |
| target x intensity | 0.59 | .447 | 0.001 |
| hand x intensity | 0.03 | .874 | < 0.001 |
| target x hand x intensity | 0.15 | .698 | < 0.001 |

157

Table A13

*ANOVA of right primary somatosensory cortex.*

| Effects | $F_{(1,44)}$ | $p_{(two\text{-}tailed)}$ | gen. $\eta^2$ |
|---|---|---|---|
| target | 37.40 | < .001 | 0.12 |
| hand | 5.40 | .025 | 0.008 |
| intensity | 0.11 | .742 | < 0.001 |
| target x hand | 0.62 | .435 | < 0.001 |
| target x intensity | 0.59 | .446 | 0.001 |
| hand x intensity | 0.21 | .645 | < 0.001 |
| target x hand x intensity | 0.002 | .967 | < 0.001 |

158

#### Supplement B

Although our behavioral data did not show a transfer of the placebo analgesia effect to empathy for pain, the ratings for both hands in the empathy condition were qualitatively more similar to the ratings for the placebo hand in the self condition. In other words, although there was no difference in pain ratings between the two hands in the empathy condition, ratings for both hands seemed to be reduced in a similar way as the first-hand ratings for the placebo hand. As a previous study by Rütgen, Seidel, Silani, et al., 2015 only observed this drop of pain empathy ratings in the placebo condition, i.e. no significant difference between self- and other-related ratings in the control condition, we aimed to further explore this “generalized” reduction in pain empathy ratings of both hands and compared ratings of the different conditions using priors derived from that study. This resulted in four Bayesian paired *t*-tests comparing pain ratings in the self/control condition with pain ratings in (a) the other/control condition, and (b) the other/placebo, as well as comparing the self/placebo condition ratings with those of (c) the other/control condition, and (d) the other/placebo condition (priors were Cohen’s *d* effect sizes from the same independent *t*-tests in Rütgen, Seidel, Silani, et al., 2015: a) 0.17, b) 0.76, c) 0.54, d) 0.05; see analysis A4 in Figure 2 in the main text).

Using these priors, we found strong evidence that the other-related pain ratings in the control condition were decreased compared to the control condition in the self (a; $BF_{10} = 2.58 \times 10^6$). This decrease was similar for the other/placebo ratings (b; $BF_{10} = 2.78 \times 10^6$). In line with this, we did not find evidence for a difference in ratings between the self/placebo and other/control conditions (c; $BF_{10} = 0.27$). Finally, similar to Rütgen, Seidel, Silani, et al., 2015, we observed comparable ratings in the placebo conditions for self and other (d; $BF_{10} = 0.85$). In other words, the placebo induction reduced the self- and other-related pain ratings to a similar extent in our study, as reported by Rütgen, Seidel, Silani, et al. (2015), but we report an additional reduction of empathy ratings for the control hand. These findings are in sharp contrast to the previous study which did not find such a decrease in the other-related control condition. In sum, although our initial predictions regarding a localized transfer to empathy were not confirmed, we found that placebo analgesia affected empathic responses in a generalized way. Specifically, we observed lower other-related pain ratings for both hands that were more similar to the self-related pain ratings of the placebo hand. We now want to discuss these results in relation to the results by Rütgen, Seidel, Silani, et al. (2015).

This mismatch of results could be explained by an inherent limitation of our within-subjects design and the effects of placebo analgesia on endogenous opioids. The opioid system generally mediates pain processing (Baumgartner et al., 2006; Karjalainen et al., 2017) and plays an important role during placebo analgesia (Büchel et al., 2014; Colloca, 2019; Eippert et al., 2009; Zubieta et al., 2005), but is also involved during empathic responses to others in pain (Rütgen et al., 2015, 2018). During placebo analgesia, two types of expectancies should be differentiated (Atlas & Wager, 2014; Geuter et al., 2017; Montgomery & Kirsch, 1996). While treatment-related expectancies relate to the overall knowledge of the context and general subjective beliefs, likely depending on sustained mechanisms such as affective shifts and tonic opioid binding (Atlas et al., 2010; Atlas & Wager, 2012), stimulus-related expectancies are hypothesized to adapt flexibly based on discrete events by depending on transient, phasic responses in systems involved in processing e.g. prediction errors (Atlas et al., 2010). For instance, Atlas & Wager (2014) provided meta-analytic evidence that treatment expectancies are significantly more likely to reduce activation in left insula than stimulus-dependent expectancies.

The within-subjects, localized placebo analgesia induction in our study could therefore have led to two distinct responses: 1) An initial tonic endorphin release over the endogenous opioid system, increasing baseline endorphin levels throughout the whole experiment. This increase, acting as a natural analgesic, might have downregulated emotional arousal and/or the affective response in general (see e.g. Benedetti & Amanzio, 2013 for a review); 2) A phasic modulation adapting flexibly depending on the location of the expected task stimulus (placebo vs. control hand, self vs. other). Such an increased tonic response in the placebo condition might have been an advantage for the previous between-subjects study who was able to induce these changes specifically in the placebo group, with the control group staying at baseline endorphin levels (Rütgen et al., 2015). This, in turn, could have eased the detection of modulation of affective responses. In our study, however, all participants underwent a placebo analgesia induction and could have experienced this tonic endorphin increase, possibly leading to a generalized down-regulated response. Our results therefore additionally provide tentative evidence for shared affective representations in the form of a generalized placebo analgesia response during empathy for pain. This idea of a primarily tonic placebo effect compared to a phasic modulation is bolstered by our Bayesian analyses showing a generalized decrease of other-related pain ratings of both hands down to the levels of the self-related placebo decrease and contradicts our initial hypothesis of a more phasic, localized response in regard to stimuli applied specifically to the placebo hand. Our within-subjects design could therefore have hindered us from observing modulation of the affective and/or somatosensory component exactly for this reason. This would then explain why we only partly replicated the behavioral, but not the fMRI results of Rütgen, Seidel, Silani, et al. (2015) regarding a placebo-induced decrease in brain regions related to the affective-motivational component for first-hand or empathic pain.

If this theory holds, however, we should also have observed a generalized/tonic response for first-hand painful experiences. Although we found evidence for a strong localized placebo effect and thus involvement of the phasic component on the behavioral level, we observed no placebo-induced decrease in brain regions related to the affective-motivational component of first-hand pain processing (except for a trend in left AI). However, this inconsistency could e.g. be explained by easier discrimination between one's own hands, i.e. higher saliency of first-hand compared to empathic experiences (see Borsook et al., 2013; Eisenberger, 2015 for critical discussions; Krahé et al., 2013; Legrain et al., 2011). These prerequisites could have allowed for a more phasic modulation in the self, while this was not or much less the case for the other, where differentiation between the hands might have demanded more complex cognitive processing.

In sum, the contribution of a "tonic placebo response" to pain processing might have had a much higher weight in our study than phasic modulatory differences between targets and hands. Our results therefore speak for an affect-modulating role of the opioid system

(Nummenmaa & Tuominen, 2017) and enhanced situational modifiability of somatosensory responses for the self compared to the other.
